# Micromotion derived fluid shear stress mediates peri-electrode gliosis through mechanosensitive ion channels

**DOI:** 10.1101/2023.01.13.523766

**Authors:** Alexandre Trotier, Enrico Bagnoli, Tomasz Walski, Judith Evers, Eugenia Pugliese, Madeleine Lowry, Michelle Kilcoyne, Una Fitzgerald, Manus Biggs

## Abstract

Clinical applications for neural implant technologies are steadily advancing. Yet, despite clinical successes, neuroelectrode-based therapies require invasive neurosurgery and can subject local soft-tissues to micro-motion induced mechanical shear, leading to the development of peri-implant scaring. This reactive glial tissue creates a physical barrier to electrical signal propagation, leading to loss of device function. Although peri-electrode gliosis is a well described contributor to neuroelectrode failure, the mechanistic basis behind the initiation and progression of glial scarring remains poorly understood.

Here, we develop an *in silico* model of electrode-induced shear stress to evaluate the evolution of the peri-electrode fluid-filled void, encompassing a solid and viscoelastic liquid/solid interface. This model was subsequently used to inform an *in vitro* parallel-plate flow model of micromotion mediated peri-electrode fluid shear stress.

Ventral mesencephalic E14 rat embryonic *in vitro* cultures exposed to physiologically relevant fluid shear exhibited upregulation of gliosis-associated proteins and the overexpression of two mechanosensitive ion channel receptors, PIEZO1 and TRPA1, confirmed *in vivo* in a neural probe induced rat glial scar model. Finally, it was shown *in vitro* that chemical inhibition/activation of PIEZO1 could exacerbate or attenuate astrocyte reactivity as induced by fluid shear stress and that this was mitochondrial dependant.

Together, our results suggests that mechanosensitive ion channels play a major role in the development of the neuroelectrode micromotion induced glial scar and that the modulation of PIEZO1 and TRPA1 through chemical agonist/antagonist may promote chronic electrode stability *in vivo*.

**Graphical abstract:** 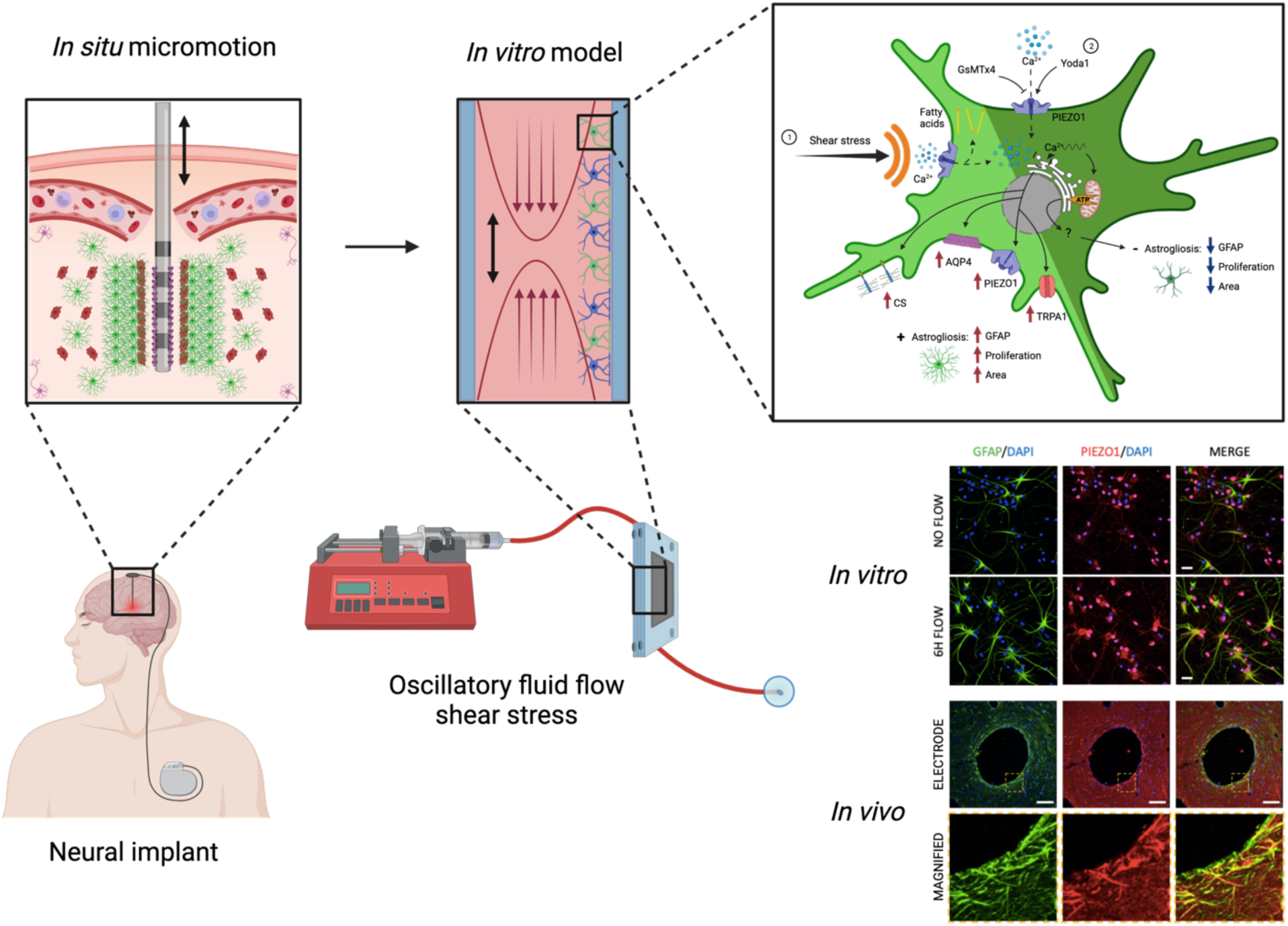

**Highlights:**

- Peri-electrode void progression is mediated by fluid flow shear stress
- Oscillatory fluid flow shear stress replicates neuroelectrode glial scarring in vitro
- Astrocyte PIEZO1 and TRPA1 are upregulated at the peri-electrode region in response to electrode micromotion
- PIEZO1 pharmaceutical activation diminishes shear stress-induced gliosis
- PIEZO1 chemical inhibition exacerbates gliosis and reduces mitochondrial functions

## 1. Introduction

Technical innovations in bioelectronic technologies have facilitated the development of smaller, smarter, and less invasive diagnostic and therapeutic devices. Current research into interfacing electronic technologies with tissues and biological system have focused on increasing the long-term performance of electrode technologies *in vivo*. The development of stable, tissue-interfacing technologies represents a paradigm shift in the treatment of chronic disease and real-time health-monitoring and an understanding of the mechanical instigators and contributors to the pro-inflammatory events which occur at the per-electrode region is critical to mitigate the detrimental effects of implant encapsulation on neural recording quality and stimulation stability [1].

Chronic neuroelectrode function is often compromised by the foreign body response, resulting in peri-electrode fibrosis and encapsulation, a process termed reactive gliosis [2–4], which can result in the complete loss of neuroprosthestic functionality. The associated inflammatory processes of gliosis are thought to be mediated principally by resident astrocytes, microglia and NG2 glia cells [1,5], which adopt a reactive and proliferative phenotype in response to localised tissue damage, leading to neuronal cell loss and the deposition of a collagen and chondroitin sulfate rich fiberous tissue [6,7].

Initially, peri-electrode inflammation is triggered by the mechanical trauma to blood vessels, capillaries and cells during implant insertion, originally proposed as the principal initiator of glial scar development [2,8–10]. Attempts to attenuate implantation-associated trauma have focused on modifying the electrode rigidity [11–15], geometry [16–19] or by coating the neural implant with anti-inflammatory chemistries to improve device integration [20–23]. However, more recent studies have shown that this initial inflammatory response to the mechanical damage of implantation is quickly resolved upon device removal [2,9]. Furthermore, a number of studies have identified a significant role of the mechanical mismatch between stiff neural probes and the soft brain composition on reactive gliosis [24,25], leading to the development of softer or mechanically adaptive neural electrodes [26–29]. In particular, a growing body of evidence shows that relative micromotion at the probe/tissue interface arising from respiration, pulsatile blood flow and physical movement [30–32], is a significant mediator of glial scarring [33–37]. These mechanical actions expose the peri-electrode tissues to cyclic shear stresses [38–44], resulting into continuous mechanically induced trauma exacerbating the inflammatory state initially primed by neuroelectrode insertion [45–48]. Critically, under acute conditions the electrode interface is unstable and the peri-electrode region is characterised by the development of a progressive fluid fill space, which appears to reach a steady state at approx. 3x the electrode diameter [33,34].

Despite insightful advances in mechanobiology, the mechanism by which cells of the CNS can sense mechanical variations and the molecular pathways which initiate gliosis are not fully understood. Recently, mechanosensitive (MS) ion channels have come to the forefront of neuromechanobiology and have been shown to undergo activation or differential expression in response to mechanical stimuli within the central nervous system (CNS) [49], with important functions in blood-brain barrier maintenance [50], neural differentiation [51–55], nociception [56–59], neural-glia communication [60–62] and glia activation [63–67]. Critically, MS channels have been observed as perturbed in neurodegenerative disorders [68] including Alzheimer’s disease [69–71], myelination disorders [72–75], migraine [76–79] and glioma [80–82].

In order to gain insight into the role of cyclic mechanical forces in the onset and evolution of gliosis, recent studies have described the development of cell culture models which mimic physiologically relevant neuroelectrode micromotions and which have demonstrated partial success in reproducing the processes of gliosis *in vitro* [83–85]. It follows that the development of pathophysiological models of the peri-implant microenvironment that replicate the glial scarring process may provide valuable tools for further understanding the mechanobiology of neural populations and promote the development of novel approaches to mitigate the foreign body response to neural implants.

In this work, we developed a novel computational model of the peri-electrode region, encompassing solid, liquid and viscoelastic elements, representing a neuroelectrode, the fluid filled peri-electrode space and the surrounding neural tissues respectively. We proceed to show that in a chronic setting, micromotion induced peri-electrode fluid shear stresses occur in the mPa range and that rat ventral mesencephalic E14 embryonic cells exposed to this level of fluid shear through a parallel-flow chamber undergo gliosis-associated cellular processes *in vitro*. It was further observed that mPa fluid shear stress induced the overexpression of two mechanosensitive ion channel receptors *in vitro*, PIEZO1 and TRPA1, which was replicated in an *in vivo* neural probe encapsulation scar model. Moreover, we show that the inhibition of PIEZO1 triggers and increases gliosis response, and hinders mitochondrial functions *in vitro*, in ventral mesencephalic cells. Conversely, chemically induced activation of the PIEZO1 receptor stalled the neuroinflammatory response of the culture when subjected to shear stress conditions. Thus, it can be hypothesised that the modulation of PIEZO1 could prevent the development of the neuroelectrode-induced foreign body reaction and enhance device integration to support long-term neural recording/stimulation application.

## 2. Methods

### 2.1. Computational modelling of neuroelectrode micromotion within the filled peri-electrode fluid space

A finite volume model (FVM) reproducing wall shear stress (WSS) at the electrode / peri-electrode fluid space / brain tissue interface was developed using ANSYS Workbench 2021 (ANSYS, Inc., Canonsburg, PA, USA). A two-dimensional parametric model of a neuroelectrode device (based on SNEX-100 electrode, Microprobes for Life Science, Gaithersburg, USA) featuring a cylindrical stepped tip was developed in SpaceClaim. The peri-electrode region filled with brain interstitial fluid was represented by a cavity between the electrode and the brain tissue **(Figure 1)**. The generated quadrilateral numerical mesh of the fluid domain consisted of 1376 cells and 1557 nodes (physics preference - CFD; solver preference - Fluent) and was validated in the Fluent / Setup module. The brain interstitial fluid density was assumed to be 1006 kg/m^3^ (based on cerebrospinal fluid density, [86]), and a viscosity range from 1.2 to 100 mPa·s was selected based on previous work [87,88]. The electrode tip was assigned material properties of stainless steel 316L, austenitic, AISI 316L, according to data compiled by Ansys Granta. The mechanical properties of brain tissues were extracted from the literature (Density 1060 kg·m^−3^, Young’s modulus 6 kPa, Poisson’s ratio 0.45) [41,89], The temperature was constant at 37 °C.

**Figure 1.**
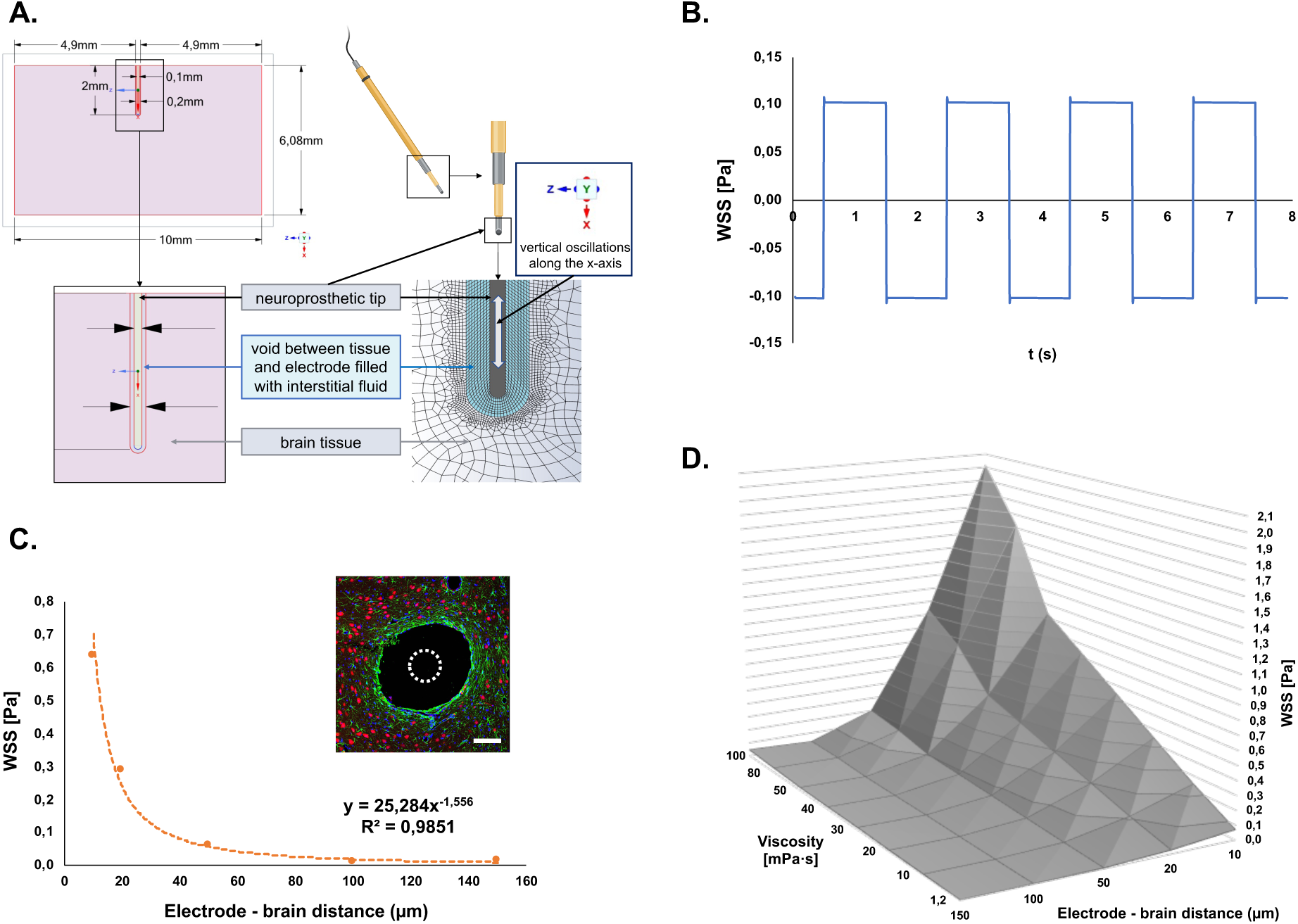
Neuroelectrode micromotion generates oscillatory interstitial fluid flow shear stresses determined by the extracellular space dimension and viscosity. Schematic of the model configuration for FVM analysis, including neural probe and mesh **(A)**. The simulated electrode movement induced oscillatory WSS at the brain tissue surface with constant frequency and amplitude **(B)**. Representative picture of an implantation tract horizontal cross-section in the STN of adult rats stained for GFAP (green), NeuN (red) and DAPI (blue) (scale bar = 100 µm), the dotted white circle symbolises the electrode position and diameter (100 µm) showing the extracellular space created around the probe. The WSS magnitude increases as the electrode – brain surface distance shortens due to cavity size reduction **(C,D)** and with the viscosity of the interstitial fluid **(D)**.

Transport equations were based on the viscous SST k - ω turbulence model and electrode micromotion was simulated by the displacement of the probe along the x-axis while the tissue remained relatively fixed. Based on previously published animal model experimental results, we applied a micromotion amplitude of 15 μm (30 μm peak-to-peak) with a frequency of 0.5 Hz arising from respiration [43,90,91]. The WSS was an average value calculated at the tissue wall located along the oscillation direction. The calculated WSS value was then used to inform the flow parameters of the *in silico and in vitro* parallel plate flow chamber (PPFC) study.s

### 2.1. Numerical 3D-simulation of oscillatory wall shear stress in the parallel plate flow chamber

To determine the consistency of fluid forces on a cell monolayer, computational fluid dynamics (CFD) characterization of the developed PPFC was performed using ANSYS Fluent **(Figure S1)**. A 3D model was reconstructed in SpaceClaim from technical sheets provided by the PPFC’s producer (Department of Mechanical & Manufacturing Engineering, School of Engineering, Trinity College Dublin, Ireland) which was further verified with chamber dimensions measurements using both digital electronic depth gauge and Vernier calliper. The generated numerical mesh consisted of 794.859 nodes and 426.604 elements (physics preference, CFD; solver preference, Fluent). The properties of the polycarbonate material used for the PPFC construction were based on the Fluent Solid Materials database (density 1200 kg/m3, specific heat 1250 J/(°C·kg), thermal conductivity 0.2 W/(°C·m)). The fluid domain consisted of DMEM/F12 - 1% FBS solution (density 1009 kg/m3, dynamic viscosity 0.930 mPa·s) [92]. A transient analysis of the viscous model at 37 °C was carried out to simulate the oscillatory fluid flow in the microchannel with a frequency of 0.5 Hz as a result of the change of the flow direction at the PPFC inlet. A volumetric flow rate in the range of 4.49 to 381 ml/min **(Table S1)** was applied.

### 2.2. Animal ethical statement

Research and animal procedures were performed in accordance with the European (EU) guidelines (2010/63/UE) and Health Products Regulatory Authority. Every effort was made to minimize animal suffering and to reduce the number of animals used.

### 2.3. Ventral mesencephalic primary cell culture from E14 rat embryos

The mesencephalon is a principal implantation site for deep-brain stimulation electrodes, and in order to assess the *in vitro* response of a clinically representative sub-population of CNS neurons ventral mesencephalic (VM) cells were extracted from E14 rat embryos of female Sprague Dawley rats according to methods described previously [12,13]. Briefly, following decapitation under anaesthesia induced by inhalation of isoflurane, embryonic sacs were extracted and placed in ice cold Hanks blank salt solution (HBSS). The embryos were then carefully removed from their sacs, ventral mesencephalons were dissected from the brains and mechanically dissociated with a pipette until obtention of a homogenous cell suspension. Cells were then seeded on poly-L-lysine-coated plain glass slides (2947-75X38, Corning®, Dublin, Ireland), where 3 divisions were previously engraved, each subdivision received 300, 000 cells. The slides were then placed in a 100 mm Petri-dish (CLS430293, Corning®, NY, US) covered by 10 mL of Dulbecco’s modified Eagle’s medium/F12 (DMEM/F12, D6421, Sigma-Aldrich, Dublin, Ireland) supplemented with 6 g/L of D-glucose, 100 µg/ml of Primocin™ (InvivoGen®, Toulouse, France), 2 mM of L-glutamine (Sigma®, Wicklow, Ireland), 10 ml/l of Fetal bovine serum (FBS) and 20 ml/l of B27® supplement (17504-044, Gibco™, NY, US), which was replaced every two days until their utilisation in different experiments.

### 2.4. Oscillatory fluid shear stimulation

Oscillatory fluid shear (OFS) was applied using a parallel-plate flow chamber (PPFC) previously designed and optimised as described by Stavenschi et al [93–95]. Briefly, a glass slide (75 x 38 x 1 mm) covered with cells was placed between two plates and an oscillatory pressure-driven fluid flow was propelled through it using 5 mL syringes (Terumo®) mounted on a programmable double syringe pump (AL-4000; World Precision Instruments®, Hertfordshire, UK). The flow rates of 22.45 mL/min and 4.49 mL/min at 0.5 Hz were used to obtain a shear stress magnitude of 0.1 and 0.5 Pa, respectively. The VM cells were lysed for protein, GAGs and RNA extraction or fixed for immunocytochemical staining after 1, 7- and 14-days post-exposition of fluid shear for 1, 4 or 6 hours **(Figure S3)**. Under static control conditions cell cultures were similarly maintained within the chambers but were not subjected to OFS.

### 2.5. Immunocytochemistry

Cells were fixed for 15 min using a 4% Paraformaldehyde solution, followed by three washes in 1X PBS and a 5-min permeabilization process using a chilled buffer containing 0.103 g/mL sucrose, 2.92 mg/mL NaCl, 0.6mg/mL, MgCl2, 4.76 mg/mL HEPES buffer and 0.1% Triton X-100 in water at pH 7.2. Non-specific targets were blocked at 37°C for 30 min using PBS containing 1% BSA. Primary antibodies (see **Table S1** for suppliers and dilutions) were incubated overnight at 4°C on a rotary shaker at 100 rpm. The next day, cells were washed 3 times with PBS-T 0.05%. Fixed samples were incubated with secondary antibodies **(Table S1)** for 1h and protected from the light at room temperature under slow shaking. Samples were then washed two times for 5 min with PBS 1X and coverslip mounted with Fluoroshield™ with DAPI (Sigma, F6057). Z-stack images were obtained at a 60-x magnification using an FV1000 Fluoview Confocal Laser Scanning Biological Microscope (Olympus®, Dublin, Ireland. All acquisition settings remained the same between cell samples allowing intensity quantification and comparison. Images were processed using ImageJ (W.RasBand, National Institute of Health, Bethesda, US).

### 2.6. Electrode implantation in vivo study

Research and animal procedures were approved by the UCD Animal Research Ethics Committee (AREC 17-22) and licences by the Health Product Regulatory Authority of Ireland (AE18982-P122).

In this study, a deep-brain stimulation model was employed, where electrodes were implanted into the subthalmic nuclei (STN) region of seven male Wistar rats of 9 weeks old and weighing approximatively 385 g (±15.4 g) at the time of surgery. Prior to surgery, rats were housed in stable pairs for a minimum of 1 week, in a controlled environment pathogen free facility, with a 12/12 hour light-dark cycle regime, access to water and standard rodent diet ad libitum, a temperature of 21.7°C (±0.017°C) and humidity of 48.8% (±0.195%). Due to possible headstage damages from the food racks, rats were fed on the ground. Cages were cleaned 3 times a week and contained paper shreds, woodchip bedding and wooden balls or sticks as enrichment.

On the day of surgery, rats were brought to the procedure room and anaesthetised using 4.5% Isoflurane in 4 L/min oxygen, the depth of anaesthesia was verified with pedal withdrawal and corneal reflex prior to 1.2-1.8% Isoflurane in 1L/min oxygen for anaesthesia maintenance. Antibiotics (Metronidazole (20mg/kg, QD) and Gentamicin (6mg/kg, QD)) and analgesia (Buprenorphine 0.015 – 0.03 mg/kg, BID) were administrated subcutaneously pre-operatively. For body temperature stability, rats were laid on a heating blanket at 37°C in the aseptically prepared surgical field throughout the entire surgery. Next, rats were placed on a Stoelting® stereotaxic frame (Dublin, Ireland) using non-rupture ear bars and local anaesthetic cream (Emla® 5% cream, AstraZeneca, Cambridge, UK) on both rat ears and frame bars. Eyes’ dryness was avoided by using a tear replacement ointment (Vidisic, Dr. Gerhard Mann, Chem.- pharm. Fabrik GmbH, Berlin, Germany). Prior to incision, a maximum of 0.5 ml of Lidocaine at 0.5% (diluted from Lidocaine 1%, Hameln Pharmaceuticals Ltd, Gloucester, UK) was injected into several areas of the skin above the skull. From between the eyes to between the ears an incision was drawn to expose the skull and visualize Bregma and Lambda. Four Stoelting® screws (1.59mm) were anchored into the skulls, 2 in the frontal bone on the left and right of the bregma and 2 in the parietal bone on the left and right of the lambda. Left and right subthalamic nucleus (STN) coordinates (location -3.6 DV and -2.5mm ML from Bregma and -7.6mm from the dura) were determined after corrections for the differences in rat skull size, when needed, and the 2 insertion holes were drilled using a Stoelting® burr until appearance of the dura. The dura was carefully sectioned, and SNEX-100 electrode (Microprobes for Life Science, Gaithersburg, USA) mounted on the stereotaxic frame and slowly descended to the STN location. Once in place, the electrode was secured to the skull using cyanoacrylic glue (Loctite, Henkel, Germany) and dental cement (Dentalon plus, Heraeus Kulzer GmbH, Hanau, Germany) was used to cover the entire skull and the anchoring screws. Finally, the skin was sutured both cranially and caudally around the cement cap using intra-dermal suture (Vycyl 4-0, Ethicon Inc., Somerville, USA).

The surgeries lasted approximatively 2 hours and the rats were returned to their home cage for recovery with a heating blanket beneath the cage and soft food for the first 18 hours. After recovery, rats were single-housed for up to 24 hours to recover and then reintroduced to their initial partner. The antibiotic course was given until 4 days post-op and the dose of analgesic was adjusted according to the observed pain level as measured using the Rat Grimace Score (RGS) [96].

### 2.7. Immunohistochemistry of brain tissue

At 8 weeks post-electrode implantation, animals were given sodium pentobarbital (∼90 mg kg−1) via the intraperitoneal (IP) cavity and once rats were unresponsive to tail/toe pinches, animals were perfused transcardially with phosphate buffered saline (PBS) followed by 10% formalin (Sigma-Aldrich, Arklow, Ireland), after administration of heparine (625 IU/rat, INNOHEP 2500 IU, LEO Pharma, Ballerup, Denmark). Rats were then decapitated, and the skulls were removed to post-fix the brain in a 4% PFA solution at 4°C for at least 24h prior removal of the probe from the brain. Fixed brains were soaked in a 15% sucrose solution overnight at 4°C, followed by a second equilibration in a 30% sucrose solution overnight at 4°C until the brain dropped to the bottom of the vial. Using a brain matrice, tissues were then finely dissected to obtain a cube containing the subthalamic nucleus and paraffin-embedded using the Thermo Scientific™ Excelsior™ ES Tissue Processor. Tissue blocks from a total of 18 animals (n=6 per control and experimental groups) were perpendicularly sectioned to the electrode tract into 10 μm thick sections using a Leica RM2135 microtome. Consecutive sections were mounted on SuperFrost Plus™ slides allowing tracking of the electrode z-axis and direct comparison between sections.

After deparaffination in a xylene bath for 2 x 5 min, tissue sections were rehydrated through increased ethanol concentration baths (100%; 100%; 90%; 70%) for 2 min each, followed by a final water bath for 5 min. Sections were then subjected to an antigen retrieval protocol, incubated in a Tris/EDTA solution (10 mM/1 mM; pH9) in a pressure cooker during 20 min. Next, tissue sections were permeabilised using a chilled buffer (0.103 g/mL sucrose, 2.92 mg/mL NaCl, 0.6mg/mL, MgCl2, 4.76 mg/mL HEPES buffer and 0.1% Triton X-100 in water; pH 7.2) for 5 min and non-specific targets were blocked for 30 min with a diluent solution containing 3% of bovine serum albumine (BSA) and 0.1% tween-20 in PBS (PBS-T). Following blocking, sections were incubated with primary antibodies to visualize either neuron nuclei (NeuN), astrocyte cytoskeleton (GFAP) or microglia cell body (Iba1) in diluent buffer at 4°C overnight (see **Table S1** for suppliers and dilutions). After 3 washes of 5 min in PBS-T, the corresponding secondary antibodies **(Table S1)** were incubated in diluent buffer for 1h at room temperature. Finally, sections were washed 3 times for 5 min with PBS, before being counterstained and coverslip mounted with Fluoroshield™ with DAPI (Sigma, F6057). The researcher performing the tissue staining remained blinded to the treatment groups.

### 2.8. Quantitative brain tissue analysis

Tissue section images were obtained at 20x magnification using an FV1000 Fluoview Confocal Laser Scanning Biological Microscope (Olympus®, Dublin, Ireland). All acquisition settings remained identical across all sections allowing intensity quantification and comparison. Images were processed using ImageJ (W.RasBand, National Institute of Health, Bethesda, US). Three to six sections were imaged and analysed per animal, resulting in 18 to 36 sections per control and experimental groups. For *in vivo* quantification with respect to the distance from the electrode insertion site, ROI boxes with a constant width of 200 μm and length of 50 μm were used every 50 μm and up to 250 μm radially outwards from the probe void. The mean intensity and percentage of the stained area was measured in each ROI box for GFAP and Iba1 stained sections and NeuN^+^ nuclei were counted using the “analyze particles” ImageJ tool. For each NeuN stained image, the distance of the 6 nearest NeuN^+^ nuclei from the electrode void was measured using the draw line tool. Finally, data were averaged for each group and plotted by mean ±SEM as a function of distance from the electrode site. The researcher performing the quantitative tissue analysis was blind to the different group identities.

### 2.9. Western blot analysis

Total protein from VM cell populations were harvested using 100 µl of RIPA buffer (R0278, Sigma-Aldrich®, Dublin, Ireland) supplemented with 1% of Protease inhibitor (Roche®, Basel, Switzerland), Phosphatase inhibitor cocktail I and III (Sigma-Aldrich®, Dublin, Ireland) and detached using a cell scraper (Sarstedt®, Wexford, Ireland). The lysate was then centrifuged for 15 min at 14000 rpm and 4°C, the supernatant collected and stored at -70°C. Sample protein concentrations were measured using Pierce™ BCA Protein Assay Kit (23227, Thermo Scientific, Waltham, US) and 4X Laemmli sample buffer containing 200 mM of DTT (11583786001, Sigma-Aldrich®, Dublin, Ireland) was added to 10 µg of protein before being denatured at 95°C for 5 min. All samples were then separated through a 10% SDS-PAGE gel at 120 V and then transferred to 0.45 µm pore size nitrocellulose blotting membranes (10600002, Amersham™ Protran™) using a Trans-Blot® Turbo™ Transfer System (Bio-Rad, Watford, UK). Membranes were blocked in 5% skimmed milk dissolved in Tris buffered saline (TBS) for 1 h, followed by incubation with 1:1000 primary antibodies dissolved in 5% milk-TBS-Tween 20 (T-TBS) overnight at 4°C. Blots were incubated with HRP-conjugated secondary antibodies at 1:10000 for 1h at room temperature (see **Table S1** for suppliers) and developed with SuperSignal™ West Pico PLUS Chemiluminescent Substrate (ThermoFisher®, Waltham, US) on CL-X Posure™ X-ray films (34091, ThermoScientific®, Waltham, US). Protein relative expression was quantified through band densitometry.

### 2.10. Protein antibody microarray

Nexterion slide H microarray slides were acquired from Schott AG (Mainz, Germany). CF™ 555, succinimidyl ester was purchased from Sigma-Aldrich® (SCJ4600022, Dublin, Ireland).

The protein-antibody array was constructed as previously described [12,97,98]. A total of 42 commercial antibodies (see **Table S2** for catalogue numbers and dilution detail) were buffer-exchanged with PBS and quantified using the Pierce® BCA Protein Assay Kit (23227, Thermo Scientific, Waltham, US). Approximately 1 nl was printed per feature on Nexterion H amine-reactive hydrogel-coated glass slides using a SciFLEXARRAYER S3 piezoelectric printer (Scienion, Berlin, Germany). The antibodies were maintained at 20°C in a 62% relative humidity environment during printing. Each microarray slide contained eight replicate subarrays, with each antibody spotted in replicates of six per subarray. After printing, slides were incubated in a humidity chamber overnight at room temperature to facilitate complete antibody-slide conjugation. Residual functional groups were deactivated by immersion in 100 mM ethanolamine in 50 mM sodium borate, pH 8.0, for 1 h at RT. Slides were washed 3X in PBS/0.05% Tween 20 (PBS-T) for 2 min, followed by 1 wash in PBS. Slides were dried by centrifugation (470 × g, 5 min) prior to storage with desiccant at 4°C until use. Protein fluorescent tagging was carried out using CF™ 555, succinimidyl ester according to manufacturer’s instructions. Briefly, the 1 mg vial of CF™ 555 was resuspended in 100 µL of DMSO and 4 µL of CF™ 555 solution was added in 50 µL of protein sample and 50 µL of boric acid. All the samples were left to incubate for 1h at RT. The excess fluorescent probe was removed through molecular weight and the buffer was exchanged with PBS (pH 7.4) using 3 kDa centrifugal filter units (UFC500396, Amicon® Ultra – 0.5mL, 3 kDa, Sigma-Aldrich®, Dublin, Ireland). Absorbance was measured at 555 and 280 nm for all tagged samples and calculations were performed according to manufacturer’s instructions using an arbitrary extinction coefficient of 100,000 and molecular mass of 100,000 SI unit, allowing quantification of relative protein concentration and label substitution efficiency.

Prior to use, the antibody microarray slides were allowed to equilibrate to room temperature for 30 min under desiccant. Fluorescently labelled protein lysates were incubated on microarrays as previously described [97,99–101] with limited light exposure throughout the process. Initially, two labelled samples were titrated (2.5 to 10 μg/ml) to determine the optimal concentration for all samples to obtain an extractable, non-saturated signal response (i.e.N1000 and 65,000 relative fluorescence units (RFU)) with low background (b500 RFU) for all samples **(Figure S6)**. In brief, 70 μl of each sample diluted to a concentration of 7.5 μg/ml in TBS-T was applied to each well of the microarray and incubated for 1 h at 23°C in the dark on a horizontal shaker (4 rpm). After incubation, slides were washed 3X for 2 min in TBS-T, 1X in TBS and then centrifuged until dry. Once dried, microarray slides were scanned immediately using a G2505 microarray scanner (Agilent Technologies, Santa Clara, CA, USA) using a 532 nm laser (5μm resolution, 90% laser power).

Microarray data extraction was performed as previously described [97,102,103]. In short, GenePix Pro v6.1.0.4 (Molecular Devices, Berkshire, UK) was used to extract raw intensity values from image files using a proprietary *.gal file which enabled the identification of 230 μm printed protein spots using adaptive diameter (70%–130%) circular alignment. The data were then exported to Excel (version 2010, Microsoft). Local background-corrected median feature intensity data (F633 median-B633) values were selected, and the median of six replicate spots per subarray was handled as a single data point for graphical and statistical analysis. Hierarchical clustering of normalized data was performed using Hierarchical Clustering Explorer v3.0 (http://www.cs.umd.edu/hcil/hce/hce3.html) using the parameters: no prefiltering, complete linkage, and Euclidean distance.

### 2.11. DNA content

DNA content was measured using a Quant-iT™ PicoGreen™ dsDNA Assay Kit (Invitrogen™, Dublin, Ireland) following manufacturer’s instruction. Briefly, 28.7 µL of DNA suspension or Lambda DNA standard (provided by the manufacturer) were added to wells with 100 µL of 1X TE buffer (10 mM Tris-HCl, 1 mM EDTA, pH 7.5) and 71.3 µL of 1X Quant-iT™ PicoGreen™ working solution. The fluorescent intensity was then read with a spectrophotometer using an excitation wavelength of 480 nm and an emission wavelength of 520 nm.

### 2.12. Quantitative real-time PCR

Total RNA from shear flow treated and static control cultures was extracted using the phase separation method. Briefly, the VM culture of 2 technical replicate slides were detached in 250 µL of TRI reagent (Sigma®) using a cell scraper (Sarstedt®, Wexford, Ireland) and were pooled together. Samples were transferred in a nuclease-free tube and stored at -80°C until RNA extraction. Later, thawed samples were vigorously shaken and vortexed followed by a 5-min room temperature incubation to ensure full cell lysis and entire protein and nucleic acid complex dissociation. Next, 100 µL of 1-Bromo-3-chloropropane (BCP) was added to each sample, vigorously shaken for 15s and left 5 min to incubate at RT before being centrifuged at 17,000g for 15 min at 4°C. The upper transparent RNA phase was carefully retrieved without disturbing the DNA phase to avoid any genomic contamination, this entire step was repeated 3 times to ensure a maximal RNA yield for each sample. Then, an equivalent volume of 2-propanol (∼200 to 300 µL) was added to the collected phase in addition of 1 µL of GlycoBlue™ (Invitrogen™) (a blue nucleic acid co-precipitant allowing a better visualization of the RNA pellet), samples were agitated and incubated for 5 min at RT or 30 min at -20°C, before centrifugation at 17,000g for 30 min at 4°C to obtain total precipitation of the RNA pellet. The supernatant was carefully discarded and nuclease-free 70% EtOH was used to wash the pellet with moderate shaking followed by centrifugation at 7,500 g for 5 min at 4°C. A second wash step was repeated with 100% EtOH. The RNA pellet was left to dry at RT for 5 min and resuspended in 14 µL of THE RNA storage solution (Invitrogen™) supplemented with 1 µL Protector RNase Inhibitor (Roche). The RNA concentration was measured using a NanoDrop 2000c Spectrophotometer (ThermoFischer®) and the purity and quality was verified using a 260/280 nm and 260/230 nm absorbance ratio, only samples with a ratio between 1.8 and 2.2 were kept for cDNA synthesis.

To avoid false amplification, the genomic DNA from each sample was eliminated prior cDNA synthesis, both steps were performed using the RT2 First Strand Kit (Qiagen®) and following manufacturer’s instructions, all steps were carried out on ice. First, 2 µL of the GE buffer was mixed with 450 ng of RNA, the volume was adjusted to 10 µL with RNase-free water and incubated at 42°C for 5 min, constituting the DNA elimination mix. Next, the RT2 First Strand Kit reverse transcription mix was prepared with 4 µL of 5X BC3 buffer, 1 µL of control P2, 2 µL of RE3 reverse transcriptase mix and 3 µL of RNase-free water. Next, 10 µL of the reverse transcription mix was added to each DNA elimination mix, tubes were gently homogenized, and incubated for cDNA synthesis at 42°C for 15 min in a Veriti Gradiant Thermal Cycler (Applied Biosystem®). The reaction ended with a temperature rise at 95°C for 5 min to inactivates the reverse transcriptase enzyme. Finally, the newly synthetized cDNA was diluted in 91 µL of RNase-free water and stored at -80°C.

Gene expression from ventral mesencephalic cell populations exposed to oscillatory and static control fluid flow conditions was pooled from 4 biological replicates and analysed via qRT-PCR using a Qiagen® RT² Profiler™ PCR Array Rat Neurogenesis and RT² Profiler™ PCR Array Rat Hippo Signalling Pathway (GeneGlobe ID : PARN-404Z & PARN-172Z, respectively). Following manufacturer’s instructions, the 384 wells of the array were filled with a 10 µL mix of nucleotide-free water, RT² SYBR Green qPCR Mastermix (Qiagen®) and samples cDNA and the quantitative RT-PCR was carried out with a LightCycler 480 (Roche). Results were quantified by a comparative Ct method and normalized using the 5 housekeeping genes included in the array.

### 2.13. Extracellular flux analysis

The oxygen consumption rate (OCR) and extracellular acidification rate (ECAR) of VM cells cultured with the PIEZO1 antagonist GsMTx4 and the PIEZO1 agonist Yoda1 was measured using the Seahorse Cell Mito Stress Test with the extracellular flux Seahorse XFp analyser (Agilent®), in order to assess the mitochondrial respiration and glycolysis, respectively [104,105]. Briefly, VM cells were extracted and seeded in PLL-coated Seahorse XF cell culture plates (25000 cells/well), after 5 days of culture growth and differentiation, cells were exposed to either 10 µM of DMSO, 10 µM of Yoda1 or 0.5 µM of GsMTx4 in culture medium or the culture medium only (control) for 14-days with the medium and drugs refreshed every 2 days. On the fourteenth day of treatment, as per manufacturer’s instructions, one hour prior to flux analysis, samples were placed at 37°C in a non-CO2 incubator and the media replaced by Seahorse Base XF medium supplemented with 1 mM sodium pyruvate, 2 mM L-glutamine and 10 mM glucose at pH 7.4 for CO2 and O2 equilibration. During this pre-incubation, the four drugs necessary to perform the Cell Mito Stress Test were loaded into the XF Sensor Cartridges with a pre-optimised concentration of 2 µM for the oligomycin, 1 µM of rotenone and antimycin A (provided already mixed by the manufacturer to be loaded and injected together) and 3 µM of carbonyl cyanide-4 (trifluoromethoxy) phenylhydrazone (FCCP). All the OCR and ECAR values were normalised by DNA content measured for each well using the Quant-iT™ PicoGreen™ dsDNA Assay Kit (Invitrogen™, Dublin, Ireland) as previously described by [106].

### 2.14. Statistical analysis

All data of this work represents at least three biological replicates for each of the test groups and the control group. Data is presented as the mean of the values ± standard error of the mean. Statistical significance was determined by One-way or Two-way ANOVA followed by Tukey post-hoc test to determine the statistical significance (p < 0.05), unless otherwise stated. All statistical analyses were performed using Minitab Express™ version 1.5.2.

## 3. Results

### 3.1. Neuroelectrode micromotion induces oscillatory millipascal fluid shear stresses to the peri-electrode tissue

Current numerical models of brain micromotion to understand and predict local tissue damage arising from implanted neuroprosthetics assume complete electrode tissue apposition for the analysis of tissue strain [26,41,89,107–109]. However, *in vivo*, neuronal death and progressive regression of nervous tissues from the electrode periphery suggest that the electrode/tissue interface is characterised by the presence of an intermediate and expanding peri-electrode interstitial fluid-filled void, that reaches equilibrium when it obtains a diameter of approximatively 3 times that of the implanted neuroelectrode, irrespective of electrical stimulation parameters [34,110]. Therefore, a more representative FVM of the electrode / extracellular space / brain tissue interface was developed to assess the magnitudes of fluid shear stress on the peri-electrode tissues, imparted through electrode micromotion within the void **(Figure 1)**.

In this model, the electrode oscillated within a fixed diameter fluid-filled cavity, with constant frequency and amplitude resulting from respiratory activity (amplitude of 30 μm peak-to-peak with a frequency of 0.5 Hz arising from respiration) **(Figure 1A,B)**. Electrode micromotion created a flow of the interstitial fluid which generates WSS at the tissue interface (**Figure 1B)** as a function of the electrode – brain tissue distance (**Figure 1C)**, and the interstitial fluid viscosity **(Figure 1D)**.

The average WSS magnitude was observed to vary from 0.0006 Pa for an electrode - brain distance of 150 μm and a viscosity value of 1.2 mPa·s, to a shear stress of 2.0167 Pa for an electrode - brain distance of 10 μm and a viscosity of 100 mPa·s **(Figure 1D)**. Specifically, when considering the interstitial fluid viscosities reported in the literature for healthy brain tissues (15-45 mPa·s) it was observed that the WSS imparted by fluid flow shear within the peri-electrode fluid space increased significantly as the electrode-tissue distance was reduced to less than 50 µm **(Figure 1C,D)**.

As the volume of the peri-electrode fluid space is observed to increase *in vivo* through dynamic radial expansion it can be inferred that a WSS > 0.1 Pa may result in astrocyte reactivity and neuronal loss, promoting expansion of the peri-electrode fluid space **(Figure 1B)**. With this hypothesis, a WSS of 0.1 Pa was chosen for the subsequent *in vitro* PPFC studies into astrocyte activation **(Figure S1 & S2)**.

Initially the capacity of the PPFC device to induce consistent millipascal fluid shear stresses on a cellular monolayer culture was assessed through CFD analysis **(Figure S1)**, confirming theoretical assumptions that in a parallel plate laminar flow system fluid flow WSS depends on the volumetric flow rate *Q*, the dynamic viscosity *μ*, and channel dimensions (width *b* and height *h*) as described by **Equation 1** [111].

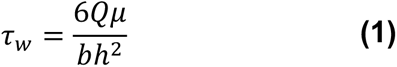

In this study, three volumetric flow rates corresponding to WSS of 0.1 Pa, 1 Pa and 10 Pa were assessed *in silico*. As the *b*/*h* ratio of the microchannel device was equal to 120, in steady conditions, 0.5 Hz oscillatory flows produced highly homogeneous WSS at 97.5% **(Figure S1C,D)** for all investigated flow regimes **(Table S1)**. Importantly, a flow rate of 4.49 ml/min provided uniform shear conditions during the flow period **(Figure S1G)** and resulted in a WSS value of 0.1 Pa **(Figure S1F)**, while the Reynolds number did not exceed 2.4.

Primary neural cells were extracted from the ventral mesencephalon of E14 rat embryos and seeded onto a glass slide five days prior to exposure to millipascal shear stress. At day 0, the glass slide was placed in the parallel flow apparatus and exposed to an oscillatory fluid flow at 0.1 Pa, 0.5 Pa and 1 Pa at 0.5 Hz for either 4 or 6 h using pulsed culture medium. Following shear stimulation, glass slides were subdivided into 3 equal pieces, and kept in culture until experiment endpoint for either PFA fixation or RNA/protein extraction **(Figure S2)**.

### 3.1. Oscillatory fluid flow shear milli-stress upregulates markers of astrogliosis and reduces neuronal viability

Increased astrocyte proliferation and GFAP expression, in association with increases in nucleus & cell area are well-established markers for astrogliosis and the glial scarring process [1,2]. To examine if the millipascal scale fluid shear stresses which arise from neuroelectrode micromotion can trigger astrocyte reactivity and neuron degeneration, we initially explored modulation of the cell morphology and cytoskeletal biochemistry in astrocyte and neuronal population in responses to oscillatory shear stress *in vitro* **(Figure 2)**. VM populations were fluorescently stained for GFAP, ß-Tubulin III and DAPI (Astrocytes, Neurons and Nuclei, respectively). Initial image analysis revealed that neuron and astrocyte populations exposed to shear stresses of 0.5 Pa and 1 Pa at 0.5 Hz were non-viable after 24 hours and these shear magnitudes were excluded from further studies (**Figure S3)**. Conversely, VM cell populations exposed to fluid flow stimulation at 0.1 Pa at 0.5 Hz for 4 or 6h, VM displayed a significant increase in the astrocyte: neuron ratio of 70% (control 19% vs treated 32%; p=0.009), after 7 days in culture following 6h of media flow **(Figure 2A)**.

**Figure 2.**
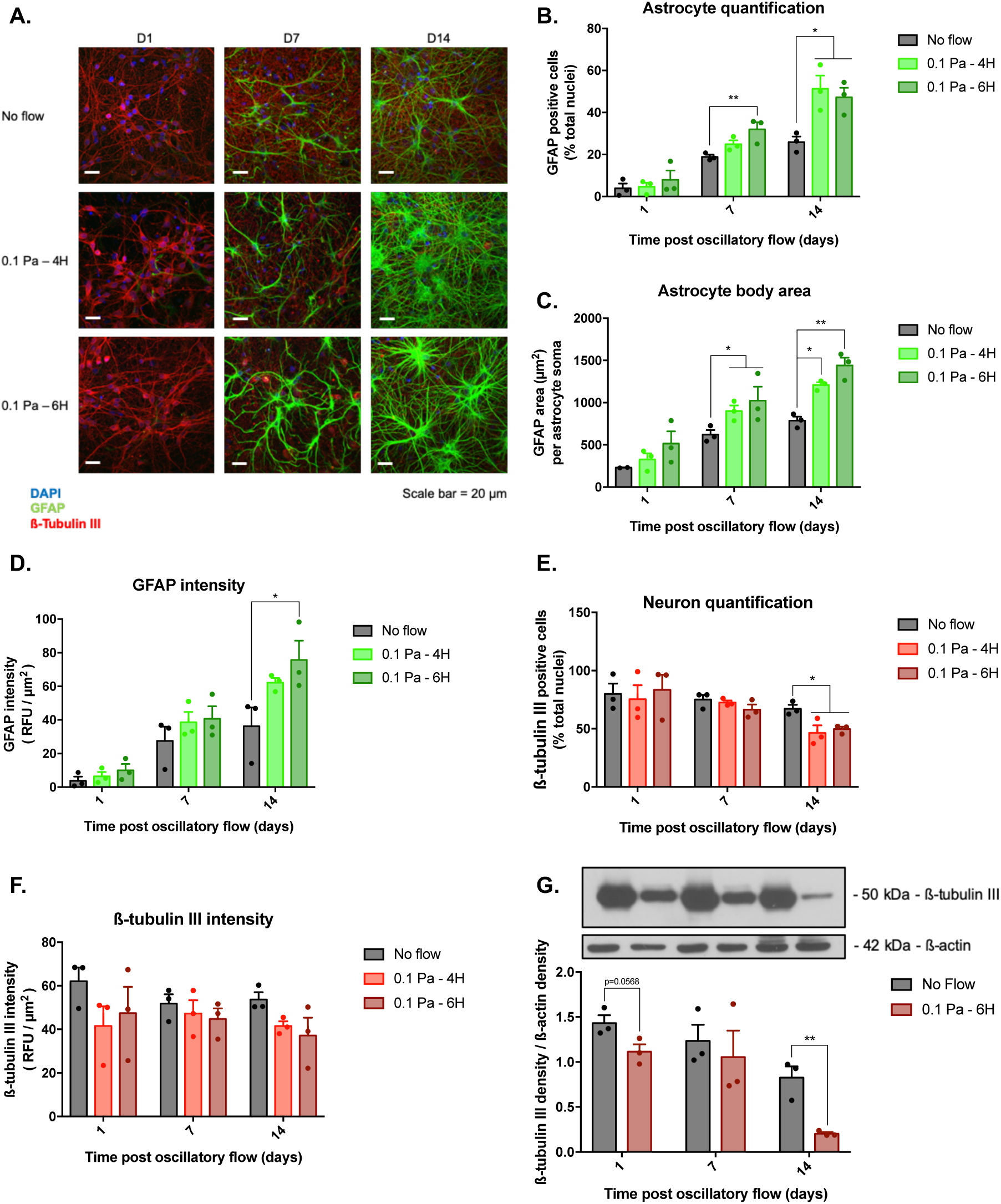
Oscillatory fluid flow shear stress promotes astrocyte reactivity and neuron death in ventral mesencephalic (VM) cells. VM cells were exposed to 0.1 Pa of fluid shear stress for 4 or 6h and immunofluorescently stained for GFAP and ß-Tubulin III after 1, 7 and 14 days **(A)** (scale bar = 20 µm; n=3). Exposure to fluid shear stress increased astrocyte proliferation by approximately 2-fold at 7 and 14 days post-flow stimulation **(B)**. The average cell area of the astrocyte was also increased by approximately 30-50% at both 7 and 14 days following oscillatory flow stimulation **(C).** Oscillatory shear stress upregulated GFAP intensity by 40 to 50% in astrocytes 14 days post stimulation **(D)**. After 14 days post oscillatory fluid flow at 0.1 Pa for 4 or 6h, a significant decrease of the total number of neurons **(E)**, and a reduction in ß-tubulin III fluorescent intensity **(F)** was observed. This reduction in ß-tubulin III was confirmed by western blot and a significant decrease of more than 65% at D14 was observed in VM cells exposed to 0.1 Pa of fluid shear stress for 6h **(G)**. Data are represented as mean ± SEM (n=3-4). One-way ANOVA with Tukey post hoc test was performed. *, ** represents a statically significant difference (p<0.05) and (p<0.01), respectively.

This effect was further observed at 14 days post-shear stress stimulation, where the astrocyte ratio was significantly increased by 98% (control 26% vs treated 51%; p=0.0242) and 82% (control 26% vs treated 47%; p=0.0482) under the 4h and 6h flow conditions, respectively **(Figure 2B)**. Shear stress also increased the astrocyte soma area, at day 7 post-flow, by 45% (control 622 µm^2^ vs treated 901 µm^2^; p= 0.0483) under the 4h flow conditions and by 65% (control 622 µm^2^ vs treated 1025 µm^2^; p= 0.0215) under the 6h flow conditions.

At 14-days post-flow stimulation, the astrocyte cell body area was also significantly increased by 54% (control 787 µm^2^ vs treated 1208 µm^2^; p=0.0241) and by 83% (control 787 µm^2^ vs treated 1441 µm^2^; p=0.0053) following 4h and 6h of fluid flow shear stress, respectively **(Figure 2C)**. Astrocytes also demonstrated an increased nuclear area of 34% (p=0.0102) and 28% (p=0.0314), following 14 days of oscillatory fluid flow stimulation for 4h and 6h, respectively **(Figure S4A)**.

Finally, following 6h of 0.1Pa shear stress, a significant >100% increase (control 36 AU vs treated 76 AU; p=0.0203) in the fluorescent intensity of GFAP was observed after 14 days **(Figure 2D)**, which was further confirmed by western blot analysis **(Figure S4B,C)**.

Conversely, 14 days following the application of 0.1 Pa shear stress, the number of neurons significantly decreased by 30% (control 67% vs treated 46%; p=0.0154) after 4h of stimulation and by 26% (control 67% vs treated 50%; p=0.0303) after 6h of stimulation, relative to cells cultured under static control conditions **(Figure 2E & S4D)**. In addition, a consistent negative trend in the overall ß-Tubulin III fluorescent intensity was observed in neuron populations stimulated under all flow conditions and at all experimental timepoints, relative to neuron populations cultured under static control conditions, in particular at 14 days post-flow exposure where a decrease of 23% (control 54 vs treated 42; n.s) and 30% (control 54 vs treated 37; n.s) was observed **(Figure 2F).** To confirm the effect of the fluid flow shear stress on ß-Tubulin III expression in VM populations, western blot analysis was carried out on cell lysates, here a reduction in ß-Tubulin III protein expression was observed in all VM populations under flow conditions. Specifically following 6h of flow, a significant reduction of 76% in ß-tubulin III protein expression (control 0.83 vs shear-exposed 0.20; p=0.0082) was observed at 14 days post-flow **(Figure 2G & S4E).**

### 3.2. Oscillatory millipascal fluid flow shear stress upregulates the expression of neuroinflammation and brain injury markers

In the CNS, traumatic and neurodegenerative events are associated with upregulations in secondary neuroinflammatory markers glycosaminoglycan chondroitin sulfate (CS) [7] and the water channel aquaporin-4 (AQP4) [112]. To determine if shear stress could similarly induce these upregulations *in vitro*, flow stimulated cultures were immunofluorescently stained for CS and AQP4 at 1, 7 and 14 days **(Figure 3A,B)**. Moreover, the proteomic expression of stimulated VM cells was further investigated using a custom protein array which targeted 8 established markers of brain injury **(Figure 3F)**.

**Figure 3.**
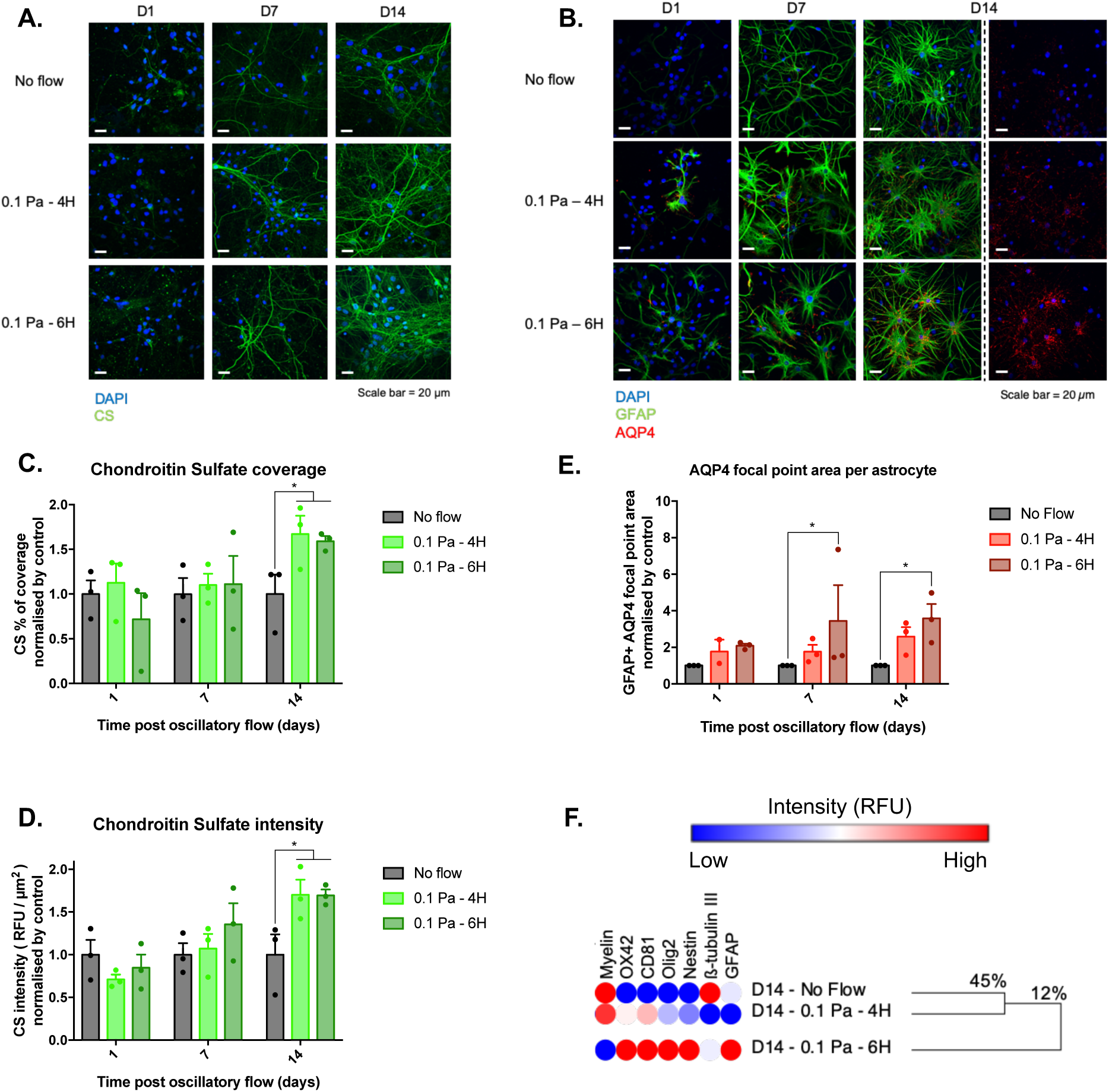
Exposing VM cells to oscillatory 0.1 Pa fluid shear stress upregulates indicators of astrogliosis, glial scarring and brain injury in vitro. The fluorescence intensity and area of Chondroitin Sulfate (CS) staining **(A,C,D)**, as well as the water channel Aquaporin-4 (AQP4) expression **(B,E)**, was significantly increased in cells exposed to flow shear stress conditions (scale bar = 20 µm; n=3). A custom antibody protein microarray was used to derive a hierarchically clustered heatmap depicting binding intensities of whole cell lysate **(H)**, (n=4). This analysis clustered together the static control condition with the 4h stimulation condition, while setting out a single group with the 6h condition, sharing only 12% similarity with the other experimental treatments. Data are represented as mean ± SEM (n=3). One-way ANOVA with Tukey post hoc test was performed. * Represents a statically significant difference of p<0.05.

It was observed that the percentage of VM cells expressing CS was increased by oscillatory fluid flow, as 14 days post-stimulation by 67% (p=0.0195) in the 4h condition and by 59% (p=0.0333) in the 6h condition **(Figure 3C)**. Upregulation in the expression of CS was also confirmed through quantification of staining fluorescent intensity which was significantly increased by 70% in both 4h and 6h treated cultures (p=0.0276 and p=0.0295, accordingly), 14 days after mechanical stimulation relative to control conditions **(Figure 3D)**.

In addition, the VM astrocyte population exhibited upregulations in AQP4 expression **(Figure 3B)**. In particular, cells subjected to 6h flow condition displayed a significant ∼2.5-fold increase at both day 7 (control 1 vs treated 3.44; p=0.0374) and 14 days (control 1 vs treated 3.58; p=0.0292) post oscillatory flow relative to VM cells cultured under static control conditions **(Figure 3E)**.

At the protein level, hierarchical analysis of the binding intensity values obtained from a custom gliosis antibody array indicated the presence of two distinct protein expression clusters 14 days post-flow **(Figure 3F)**. The first, including the no-flow control and the 4h treatment condition, displayed 45% of similarity. While, the second cluster, including only the 6h flow condition, presented only 12% of similarity with the no-flow control and the 4h treatment conditions.

The microarray analysis confirmed an increase in GFAP protein synthesis in VM cells subjected to 6h flow conditions, and a decrease in the synthesis of ß-Tubulin III in cell populations subjected to 4 and 6 hours of fluid flow shear stress. Furthermore, astrocyte reactivity was asserted by a significant upregulation in the synthesis of Nestin, while downregulation of Myelin protein further indicated neuronal loss. Finally, the protein microarray revealed the overexpression of OX42 and CD81, proteins that become upregulated in activated microglia, as well as an increase in expression of Olig2, a protein shown to have a neuro-protective role following brain injury. Together, these findings provide further evidence that the application of oscillatory fluid flow shear stress on VM cells is able to recapitulate the cellular processes of reactive gliosis *in vitro*.

### 3.3. Oscillatory millipascal fluid shear stress promotes a pro-inflammatory phenotype, neurodegeneration and modulates mechanotransduction processes

To further explore the potential influence of millipascal shear stress on neural cell function, genomic expression profiles of VM cultures exposed to shear flow and static control conditions were analysed using commercially available PCR arrays targeting 84 specific neurogenesis-related genes and 84 genes belonging to the hippo pathway family **(Figure 4A-G)**. The 18 most upregulated **(Figure 4B)** and 14 most downregulated **(Figure 4C)** genes were identified using a cut-off value of Z>2-fold or Z<-2-fold regulation (Z = flow vs static conditions).

**Figure 4.**
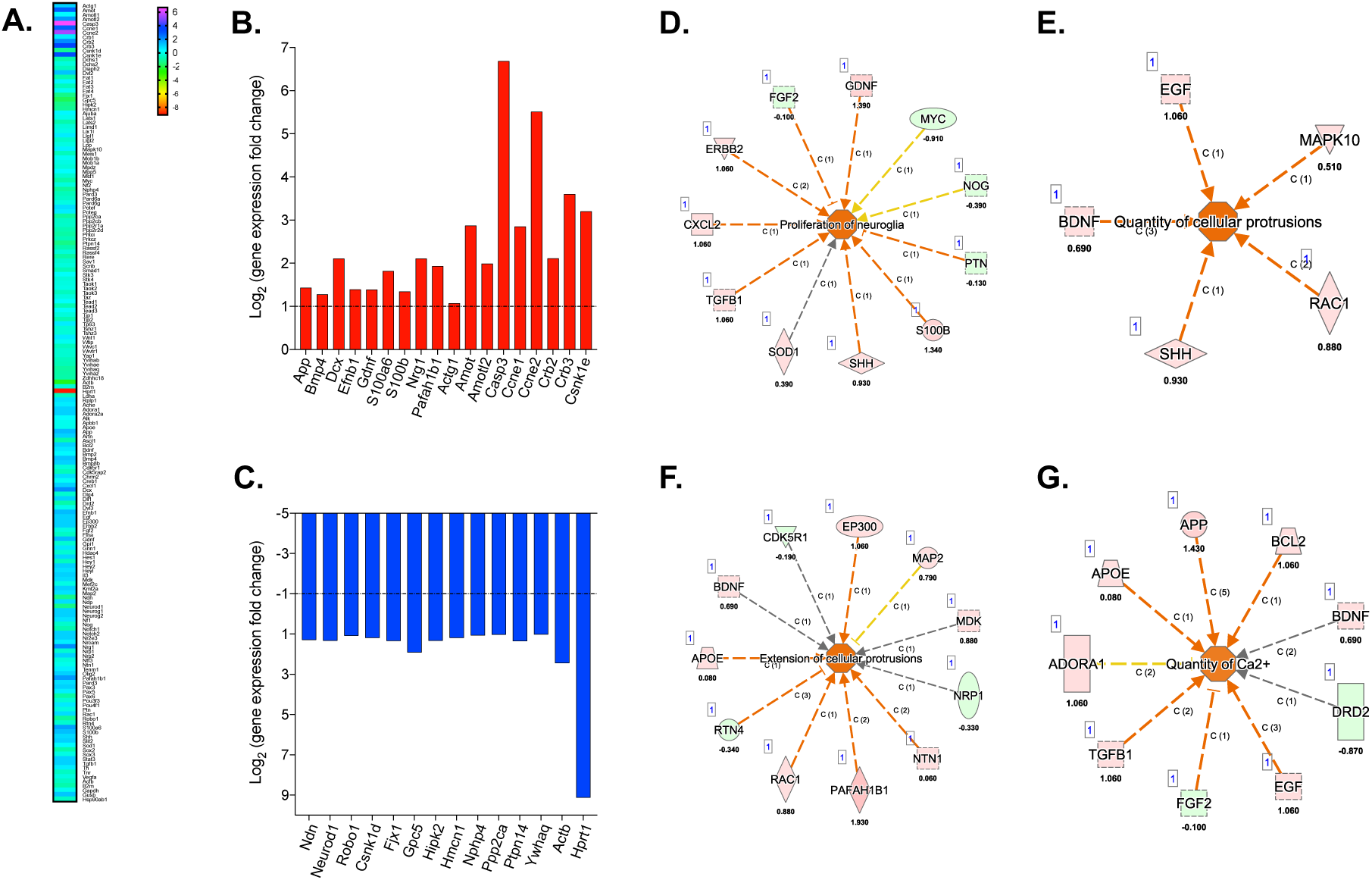
Exposing VM cells to oscillatory 0.1 Pa fluid shear stress induces glial cell proliferation and growth, neurodegeneration and modulates multiple functional pathways. The genomic expression of VM cells following 6h of flow shear stress in vitro was assessed using qPCR arrays targeting 168 genes related to neurogenesis and the hippo pathway **(A)**. The 18 most upregulated **(B)** and 14 most downregulated **(C)** genes related to astrocyte reactivity, neurodegeneration and brain injury are displayed in red and blue, respectively (Log^2^ fold regulation; dotted line representing the cut-off of ≤2-fold or ≥-2-fold regulation, respectively; n=4 pooled biological replicates). Qiagen Ingenuity Pathway Analysis (IPA) software predicted the activation (Z>2) of 4 biological function networks: Proliferation of Neuroglia **(D)**, increased quantity and extension of cellular protrusions **(E,F)** and increased quantity of Ca^2+^ **(G)** (prediction legend in **Figure S6**).

The most significantly upregulated cell-cycle associated gene in response to fluid shear stress was the apoptosis-related gene Casp3 with a 102.54-fold upregulation relative to VM populations cultured under static control conditions [113–115]. Amyloid-ß precursor protein (App), associated with neuronal death also exhibited a 2.69-fold upregulation in VM populations cultured under shear-flow conditions [116,117]. The two members of the S100 calcium-binding protein family involved in astrocyte proliferative processes, S100a6 and S100b also underwent upregulations of 3.52 and 2.53-fold, respectively [118–121]. CCNE1 and CCNE2, genes of the cyclin family, reported to play a role in cell cycle and mitotic events, showed a positive regulation of 7.19 and 45.57-fold, respectively [122]. Similarly, casein kinase I isoform epsilon gene (CSNK1E), involved in DNA replication and repair, as well as circadian regulation exhibited upregulated expression of 9.19-fold when compared to VM populations cultured under static control conditions [123].

The most downregulated cell-cycle associated gene induced by oscillatory fluid flow stimulation was HPRT1 (Hypoxanthine Phosphoribosyltransferase 1), with a 556.41-fold downregulation, a gene linked to DNA and RNA production through purine recycling [124]. Similarly, Fjx1 (Four-Jointed Box Kinase 1) and Gpc5 (Glypican 5), both involved in the control of cellular growth, division and differentiation were also downregulated by 2.51-fold and 3.75-fold respectively [125–127]. Likewise, PPP2CA, encoding the phosphatase 2A catalytic subunit and responsible for the negative control of cell proliferation and growth was also downregulated by 2.03-fold [128].

With respect to neurogenesis, the Neuregulin 1 gene, NRG1, involved in growth and differentiation of both neurons and glial cells, displayed a 4.3-fold upregulation in expression [129,130], along with the Platelet-activating factor acetylhydrolase IB subunit alpha, PAFAH1B1, a marker of neurogenesis, which exhibited a 3.81-fold upregulation [131]. Critically, Neuronal differentiation 1 (Neurod1), a transcriptional activator responsible for neuronal morphogenesis and maintenance, underwent a 2.50 downregulation in expression [132,133]. The expression of Glial Cell-Derived Neurotrophic Factor, GDNF, a promoter of neuron growth and survival underwent a 2.61-fold upregulation [134,135]. Concurrently, Bone Morphogenetic Protein 4 (BMP4), also displayed a 2.42-fold upregulation in expression, and has been shown to repress neurogenesis *in vitro* [136,137]. Finally, Angiomotin (AMOT) and angiomotin-like 2 (AMOTL2), modulators of the Hippo signalling pathway, exhibited 7.31 and 3.97-fold upregulations, respectively, in cells exposed to shear stimulation relative to cells cultured under static conditions [138].

With respect to cell cytoskeleton and motility, the gene Gamma-actin, ACTG1 showed a 2.10-fold upregulation [139]. Similarly, doublecortin, DCX, a known cytoskeletal marker of neurogenesis also exhibited a 4.3-fold increase in expression [140]. The Ephrin B1 gene, EFNB1, which also plays a role in cell adhesion also displayed a positive regulation of 2.62-fold [141]. Conversely, Roundabout guidance receptor (Robo1), an axon guidance and cell adhesion receptor, demonstrated a 2.10-fold downregulation [142,143]. Furthermore, the actin beta (ACTB) and Hemicentin 1 (HMCN1), linked to cell structural integrity and motility along with adhesion and mechanotransduciton, also demonstrated a 5.41-fold and 2.27-fold negative regulation in expression, respectively [139,144]. Interestingly, fluid-shear stress also induced a 2.27-fold downregulation of the Casein Kinase 1 Delta gene (CSNK1D), indicative of Yap1 activation, a downstream actor of the “mechanosensing” Hippo signalling pathway [123]. Moreover, the Nephrocystin 4 gene (NPHP4) and Protein Tyrosine Phosphatase Non-Receptor Type 14 (PTPN14), both act as negative regulators of the Hippo pathway by inhibiting TAZ and YAP and were downregulated by 2.27- and 2.54-fold, respectively [145–147].

Next, Ingenuity Pathways Analysis (IPA®) software was used to predict the activation or inhibition of biological functions, by creating functional networks using the differential gene expression between neural populations exposed to fluid shear or to static conditions. The IPA software predicted 6 significantly modified biological functions (Activated Z>2, Inhibited Z<-2), 14 days after VM exposure to 6 hours of 0.1 Pa of shear stress. Specifically, the upregulated expression of S100B, SHH, SOD1, TGFB1, CXCL2, ERBB2 and GDNF together with downregulated expression of PTN and FGF2 led to the predicted activation of “the proliferation of neuroglia” biological function **(Figure 4D)**. In addition, grouping of the 5 positively modulated genes EGF, MAPK10, BDNF, RAC1 and SHH predicted the activation of the “quantity of cellular protrusions” function **(Figure 4E)**, along with the predicted activation of the “extension of cellular protrusions” function due to the upregulation of EP300, NTN1, PAFAH1B1, RAC1, APOE and the downregulation of RTN4 **(Figure 4F)**. Moreover, the positive regulation of APP, APOE, BCL2, TGFB1 and EGF led to a predicted activation of the “Increased quantity of Ca^2+^” function **(Figure 4G)**.

Interestingly, two related biological functions, despite a Z-score under the threshold of significance, showed a Z>1.5 predicting the activation of the “release of fatty acid” function **(Figure S6A)** and a Z<-1.5 which predicts an inactivation of the “transport of monosaccharide” biological function **(Figure S6B)**.

Finally, the pathway identifier tool, comparing the transcriptome expression of neural populations exposed to 6 hours of flow relative to static control conditions suggested a significant upregulation of the Neuroinflammation pathway and biological functions related to proliferation and metabolism **(Figure S6C)**. Specifically, this pathway displayed a predicted activation of biological roles such as astrogliosis, neurons damage, neurofibrillary tangles, Aß generation, apoptosis, oxidative stress, ROS production, BBB disruption and Ca^2+^ overload.

### 3.4. Oscillatory millipascal fluid shear stress promotes the upregulation of mechanosensitive ion channels

Transmembrane mechanosensitive ion channels such as members of the PIEZO or TRP families have been shown to play a role in mediating cellular responses to physicomechanical stimuli through intracellular signalling cascades under both physiological and pathophysiological conditions [52,68]. To determine whether shear stress induced astrocyte reactivity was triggered by ion channel mechanoreception, an in-house protein microarray was developed to assay the expression of known mechanically gated ion channels in primary VM populations subjected to millipascal shear stress relative to static control conditions. The two most upregulated receptors were subsequently further investigated *in vitro* using immunochemistry **(Figure 5)**.

**Figure 5.**
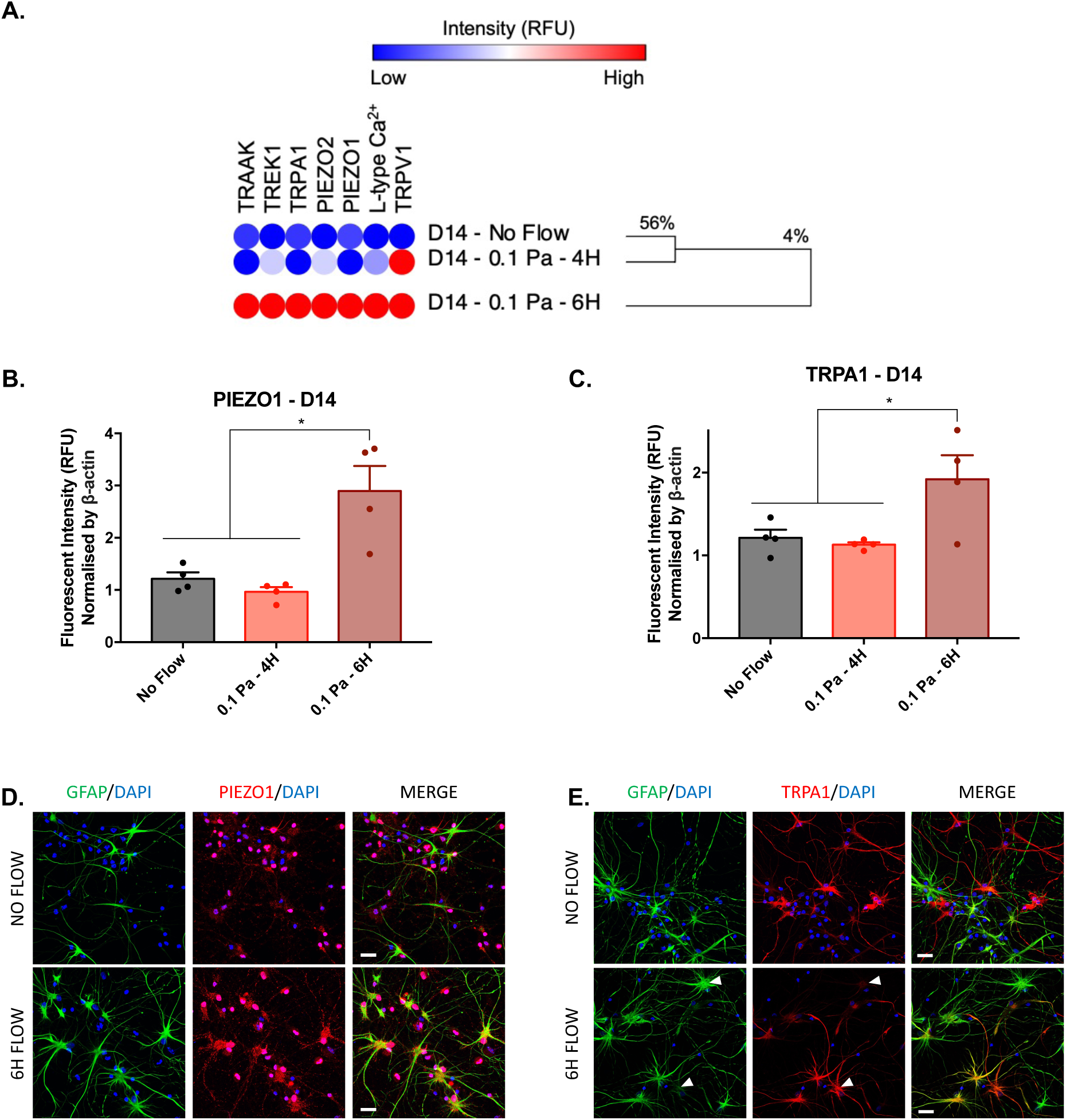
The mechanosensitive ion channel proteins PIEZO1 and TRPA1 are upregulated in response to millipascal fluid flow shear stress. A custom ion-channel antibody microarray was used to generate a hierarchically clustered heatmap depicting binding intensities using complete linkage and Euclidean distances of scale normalized by ß-actin **(A)**, (n=4). Increased expression of all ion channels 14 days after exposure to a 0.1 Pa shear flow for 6 hours was observed. Quantitative analysis of antibody microarray fluorescence intensity of PIEZO1 **(B)** and TRPA1 **(C)** 14 days after exposure to a 0.1 Pa shear flow for 6 hours. These data were further confirmed using immunocytochemistry of DAPI (blue), GFAP (green) and PIEZO1 (red) **(D)** or TRPA1 (red) **(E)**, (scale bar = 20 µm). Data are represented as mean ± SEM (n=4). One-way ANOVA with Tukey post hoc test was performed. * Represents a statically significant difference of p<0.05.

Intriguingly, *all* tested mechano-receptor proteins were upregulated following 4 h of 0.1 Pa flow shear stress, an effect which was exacerbated by increasing the stimulation time to 6h **(Figure 5A)**. Indeed, Euclidian cluster analysis using complete linkage returned two principal clusters 14 days post-stimulation: Cluster 1 contained the 4h shear flow group and the static conditions which exhibit 75% similarity and cluster 2, represented the 6h flow condition and demonstrated 7% similarity with the 4h shear flow group and the static condition cluster **(Figure 5A)**.

Among the assayed ion channels, two members of the transient receptor potential (TRP) cation channel family, the Vanilloid Receptor 1 (TRPV1) and Ankyrin 1 (TRPA1), which are known to be involved in mechanical, temperature and chemical sensing, were upregulated following shear stress stimulation. In particular, TRPA1, which has been recognised to act as calcium influx regulator in astrocytes [148,149], exhibited a significant 59% increase (control 1.21 vs treated 1.92; p=0.0492) in expression after 6h of 0.1 Pa shear stress, relative to cells cultured under static control conditions **(Figure 5C)**.

Moreover, PIEZO1 and PIEZO2 were also overexpressed following 0.1 Pa oscillatory flow shear stress, with greater expression following 6h **(Figure 5A).** Interestingly, PIEZO1 overexpression has been observed to occur in neurodegenerative disorders including Alzheimer’s disease or multiple sclerosis [70,150]. In this study PIEZO1 displayed a significant ∼2.4-fold increase (control 1.22 vs treated 2.89; p=0.0304) in expression relative to cells cultured under static control conditions **(Figure 5B)**.

PIEZO1 overexpression in VM populations exposed to fluid flow conditions was subsequently confirmed by immunocytochemistry which confirmed the presence of relatively large reactive astrocytes with a high GFAP intensity accompanied by a marked increase in PIEZO1 receptor staining in both the astrocytic somas and cell neurites **(Figure 5D)**. Interestingly, upregulation of the TRPA1 receptor intensity appeared to negatively correlate with GFAP intensity in astrocyte populations **(Figure 5E)**.

### 3.5. Neuroelectrode implantation triggers astrocyte overexpression of PIEZO1 and TRPA1 *In vivo*

To confirm the relationship between neuroelectrode micromotion induced fluid shear stress on glial scarring and astrocyte expression of PIEZO1 and TRPA1, we developed a *in vivo* rat model of glial scarring by implanting commercially available neuroelectrodes in the subthalamic nuclei. Probes were either removed immediately after insertion to create a stab wound or retained in place for 8 weeks to generate a neuroelectrode-induced glial scar **(Figure 6 & Figure S7)**.

**Figure 6.**
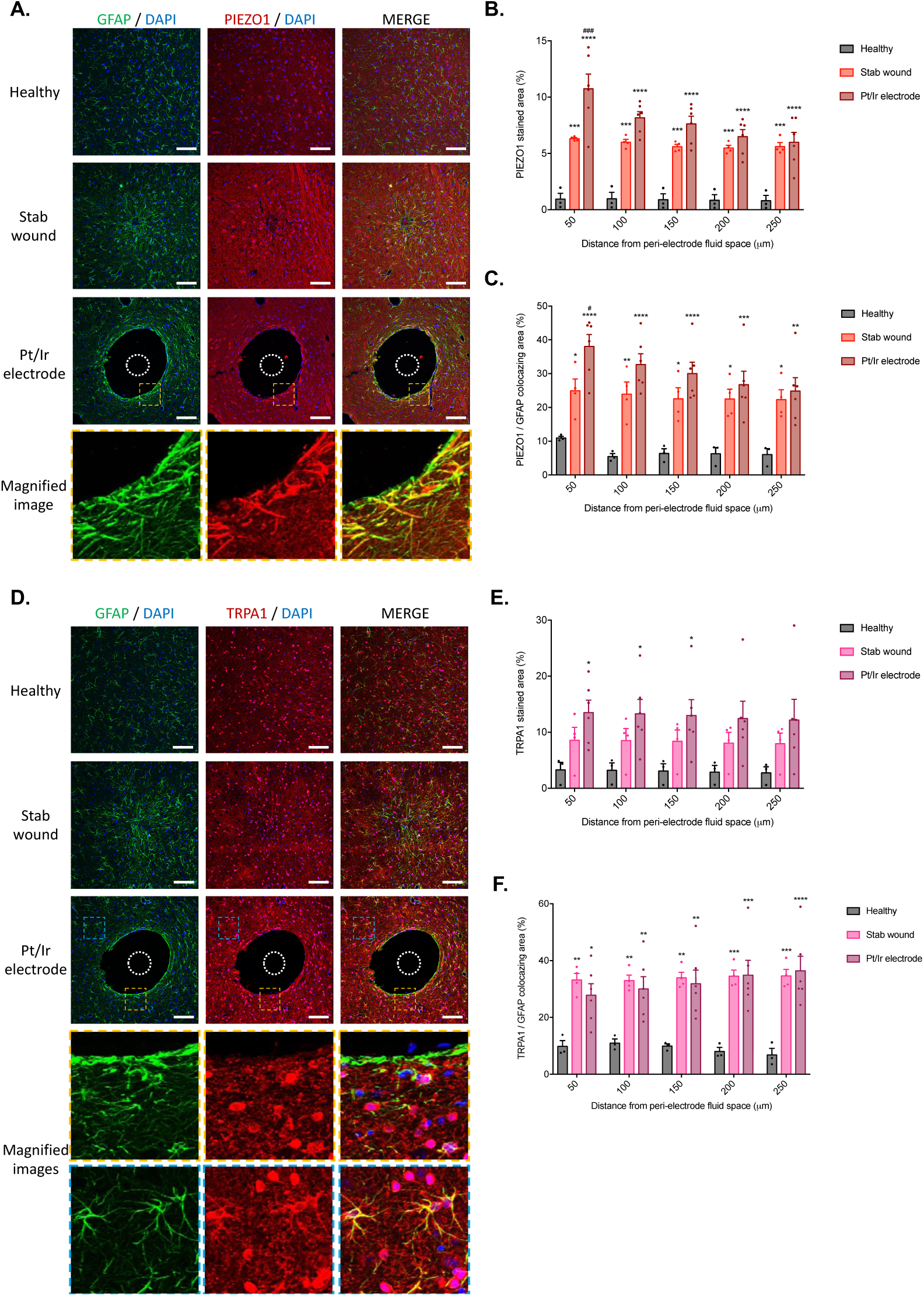
PIEZO1 and TRPA1 receptors are upregulated in vivo in peri-electrode astrogliosis. To verify the prediction of the in vitro model, Pt/Ir electrodes were implanted in the rat subthalamic nucleus and either immediately removed (stab wound) or retained in situ for 8 weeks (Pt/Ir electrode). Representative Immunohistochemistry images of non-implanted healthy control tissue, an electrode stab injury and Pt/Ir electrode-implanted brain tissue, stained in green for GFAP, red for PIEZO1 and blue for DAPI, yellow dash line depicts a zoomed region, the white dashed circle represents the electrode position (scale bar = 100 µm) **(A)**. Quantification of the PIEZO1 stained area **(B)** and colocalization of PIEZO1 to GFAP staining **(C)** relative to healthy control tissue. Representative Immunohistochemistry images of non-implanted healthy control tissue, an electrode stab injury and Pt/Ir electrode-implanted brain tissue, stained in green for GFAP, red for TRPA1 and blue for DAPI, yellow or blue dash line depicts a zoomed region, the white dashed circle represents the electrode position (scale bar = 100 µm) **(D)**. Quantification of the TRPA1 stained area **(E)** and colocalization of TRPA1 to GFAP staining **(F)** relative to healthy control tissue. Data are represented as mean ± SEM (n=3-6). Two-way ANOVA with Tukey post hoc test was performed. *, **, ***, **** represents a statically significant difference versus the healthy control and #, ##, ###, #### versus the stab wound condition, (p<0.05), (p<0.01), (p<0.001) and (p<0.0001), respectively.

Notably, a peri-electrode void of approx. 3x the electrode diameter was observed only in the *in situ* implantation group, as reported extensively [26,27,34,35]. We next assessed astrocyte and microglia aggregation and neuronal loss (well described indicators of gliosis [1,2]) at the peri-electrode region to validate the robustness of our model. Indeed, in both the stab injury and neuroelectrode insertion conditions, the GFAP stained area exhibited a significant increase of ∼70% and ∼80% respectively (p=0,0004 & p<0.0001) 50 μm away from the insertion site, and an increase of ∼56% and ∼59% (n.s & p=0.0165), respectively, 250 μm from the insertion relative to healthy control tissues **(Figure S7B)**. Similarly, electrode insertion led to a ∼63% increase in peri-electrode Iba1 staining in the stab wound group and ∼78% increase in the Pt/Ir electrode in situ group (n.s & p=0.0001, respectively) in the first 50 μm from the implantation site, relative to the healthy tissue controls group. This effect was sustained up to 150 μm away from the peri-implant fluid space and a significant ∼65% increase in microglia staining (p=0.0499) was noted for the platinum electrode in situ group **(Figure S7C)**.

Conversely, a ∼76% (n.s) decrease in peri-electrode neuron density was observed in the stab injury group and a significant decrease of ∼93% (p=0.0314) in the in-situ Pt/Ir electrode group 50 μm away from the peri-electrode fluid space, relative to healthy control tissues. Neuronal loss was also present at a distance of 250 μm from the peri-electrode fluid space and significant reductions in neuron density of ∼55% and 67% (p<0.0001 & p<0.0001) were observed, respectively **(Figure S7D)**.

The expression of the mechanosensitive ion channels PIEZO1 and TRPA1 in both the stab wound and the in situ peri-electrode glia scar was also assessed *in vivo* relative to healthy control tissue via analysis of staining area and the colocalization of PIEZO1 and TRPA1 with the astrocytic marker GFAP **(Figure 6A,D)**. Both *in situ* electrode and stab wound groups presented a marked and significant increase in staining area of PIEZO1, specifically, an increase of ∼85% (p=0.0001) for the stab injury and ∼91% (p<0.0001) for the *in situ* electrode group was observed at 50 μm from the peri-electrode fluid space, which persisted as an ∼86% increase in area (p=0.0005 & p<0.0001) at 250 μm from the implantation site, for both experimental groups **(Figure 6B)**.

Notably, PIEZO1/GFAP colocalization was observed to follow a similar trend when compared to healthy control tissues, and significantly greater colocalization was measured up to a distance of 250 μm from the peri-electrode fluid space in both experimental groups. PIEZO1/GFAP colocalization was observed to increase from ∼56% (p=0.0392) for the stab wound and ∼71% (p<0.0001) for the electrode group, 50 μm from the electrode interface, to ∼72% (p=0.0140) and ∼75% (p=0.0017), respectively, 250 μm from the peri-electrode fluid space **(Figure 6C)**.

Similarly, a significant ∼75% increase in the total stained area of TRPA1 was observed in the Pt/Ir electrode in situ group at 50 μm (p=0.0341), 100 μm (p=0.0374) and 150 μm (p=0.0418) from the insertion site relative to control healthy tissues **(Figure 6E)**. Interestingly, TRPA1/GFAP colocalization demonstrated a consistent increase as a function of distance in both experimental groups, with a ∼70% (p=0.0023) and ∼65% (p=0.0123) colocalization increase respectively within the first 50 μm, which increased to ∼80% (p=0.0003 & p<0.0001, respectively) colocalization 250 μm from the implantation site **(Figure 6F)**.

### 3.6. PIEZO1 inhibition triggers or aggravates astrogliosis, while its overactivation hinders the glial scarring process

To further understand the role played by PIEZO1 in the development of peri-electrode gliosis and to explore the potential for ion channel inhibitors or activators in anti-gliosis therapy, static and dynamic (6h, 0.1 Pa) VM cell cultures were supplemented with GsMTx4 and Yoda1, established chemical antagonists and agonists of PIEZO1 respectively for 14 days. Astrocyte and neuron density were subsequently assessed by fluorescent labelling of GFAP and β-tubulin III, respectively **(Figure 7)**. Both inhibition and activation of PIEZO1 significantly modulated astrocyte and neuron density relative to control culture conditions, effects which were further exacerbated in VM populations exposed to flow shear-stress conditions **(Figure 7A)**.

**Figure 7.**
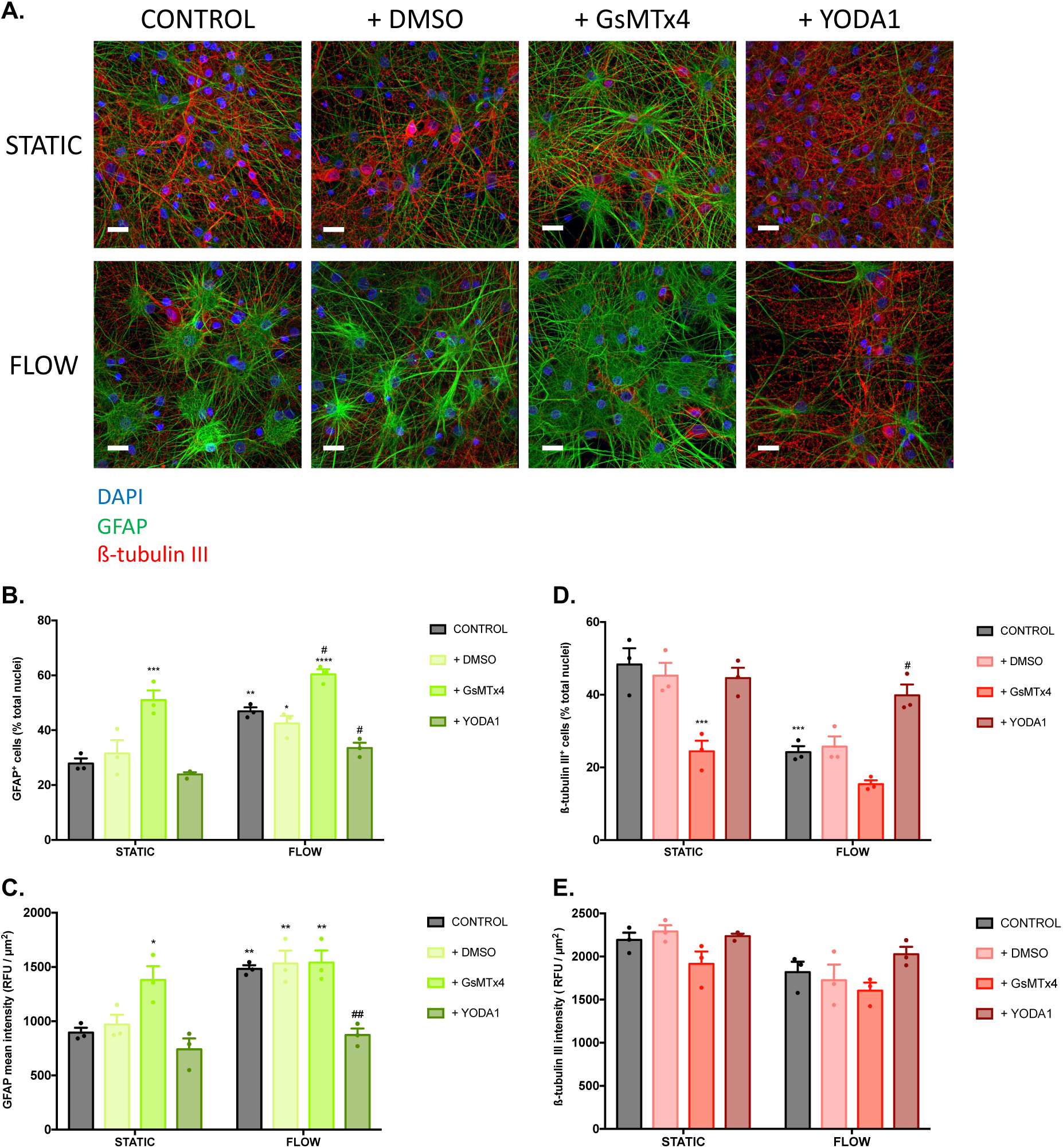
PIEZO1 inhibition promotes astrocyte reactivity and neurodegeneration while PIEZO1 activation rescues gliosis triggered by shear stress. To evaluate the action of the PIEZO1 chemical agonist Yoda1 and the PIEZO1 antagonist GsMTx4 on astrocyte reactivity and neuron viability, VM, populations were exposed to flow shear stress (0.1 Pa, 6h) and supplemented with 10 µM of Yoda1/DMSO or 0.5 µM of GsMTx4 for 14 days and stained for GFAP and ß-Tubulin III **(A)**, (scale bar = 20 µm; n=3). Quantitative analysis of GFAP positive cell number in response to Yoda1, GsMTx4 and control DMSO under flow and static culture conditions **(B).** Quantification of GFAP intensity in astrocyte populations in response to Yoda1, GsMTx4 and control DMSO under flow and static culture conditions **(C)**. Quantitative analysis of ß-Tubulin III positive cell number in response to Yoda1, GsMTx4 and control DMSO under flow and static culture conditions (**D)**, Quantitative analysis of ß-Tubulin III intensity of VM cells in response to Yoda1, GsMTx4 and control DMSO under flow and static culture conditions **(E)**. Data are represented as mean ± SEM (n=3-4). Two-way ANOVA with Tukey post hoc test was performed. *, **, ***, **** represents a statically significant difference versus the static control and #, ##, ###, #### versus the flow control, (p<0.05), (p<0.01), (p<0.001) and (p<0.0001), respectively.

Quantification of GFAP^+^ cells in a mixed VM culture under static culture conditions revealed a significant ∼45% increase in cell number (p=0.0003) following exposure to the PIEZO1 inhibitor GsMTx4, similar to the observed increase in astrocyte number resulting from culture under oscillatory fluid flow (p=0.0022). Furthermore, combining fluid shear stress stimulation with inhibition of PIEZO1 through GsMTx4 exposure amplified further the astrocyte presence to ∼54% (p<0.0001) and ∼22% (p=0.0382) significant increase relative to astrocyte presence under flow alone and static control conditions, respectively. Interestingly, activation of PIEZO1 using the chemical agonist Yoda1, inhibited the astrocyte proliferative effects induced by fluid shear stress by significantly reducing the astrocyte population by ∼29% (p=0.0395) relative to cells cultured under flow only conditions, indeed no significant differences were observed in astrocyte number in VM populations exposed to Yoda treatment under flow conditions relative to static control conditions **(Figure 7B)**.

Similarly, the analysis of mean GFAP intensity revealed a significant increase of ∼35% (p=0.0298) in cells cultured under static conditions and exposed to the PIEZO1 inhibitor GsMTx4 relative to cells cultured under static control conditions alone. GFAP intensity was increased to ∼40% (p=0.0065) when GsMTx4 supplementation was coupled with oscillatory flow stimulation relative to static control conditions. Conversely, Yoda1 supplementation of VM cells cultured under fluid flow conditions significantly decreased astrocyte GFAP intensity by ∼41% (p=0.0046) relative to the flow control condition, returning the mean GFAP intensity to that of astrocytes cultured under static control conditions **(Figure 7C)**.

In a similar manner, the number of β-tubulin III^+^ cells was quantified in response to experimental flow conditions and PIEZO1 inhibition/activation. Here, the addition of GsMTx4 to the static cultures led to a significant ∼49% (p=0.0006) reduction in the neuronal population density relative to cells cultured under static control conditions, comparable to VM cells cultured under fluid shear stress conditions alone. Critically, exposure to the PIEZO1 antagonist GsMTx4 induced a further (insignificant) ∼20% decrease in the presence of β-tubulin III positive cells relative to cells cultured under experimental flow conditions.

Conversely, exposure to the PIEZO1 agonist Yoda1 significantly protected against the loss of neurons by shear stress conditions, inducing a ∼39% (p=0.0289) increase in the number of β-tubulin III positive cells relative to cells cultured under fluid flow conditions, resulting in a comparable number to that of cells cultured under static control conditions **(Figure 7D)**. Similarly, analysis of β-tubulin III mean intensity indicated a ∼13% (n.s) reduction with GsMTx4 supplementation of cells cultured under static conditions and a ∼12% (n.s) diminution for cells under flow conditions, relative to cells cultured under static conditions alone and flow conditions alone, respectively **(Figure 7E).**

### 3.7. PIEZO1 inhibition of neural populations reduces mitochondrial oxygen consumption rate and extracellular acidification rate through glycolysis impairment

To probe the impact of PIEZO1 modulation on neural metabolic activity, we assessed the effects of its chemical activation and inhibition on mitochondrial oxygen consumption rate (OCR) and extracellular acidification rate (ECAR) *in vitro*. Naïve VM cells were exposed to the PIEZO1 agonist Yoda1 and antagonist GsMTx4 for 14 days in culture prior to extracellular flux analysis **(Figure S8A)**. Overall, it was observed that GsMTx4 supplementation negatively affected both OCR and ECAR suggesting an effect on respiration and glycolysis. Conversely, Yoda1 supplementation did not significantly modify OCR or ECAR in VM populations **(Figure 8A,B)**.

**Figure 8.**
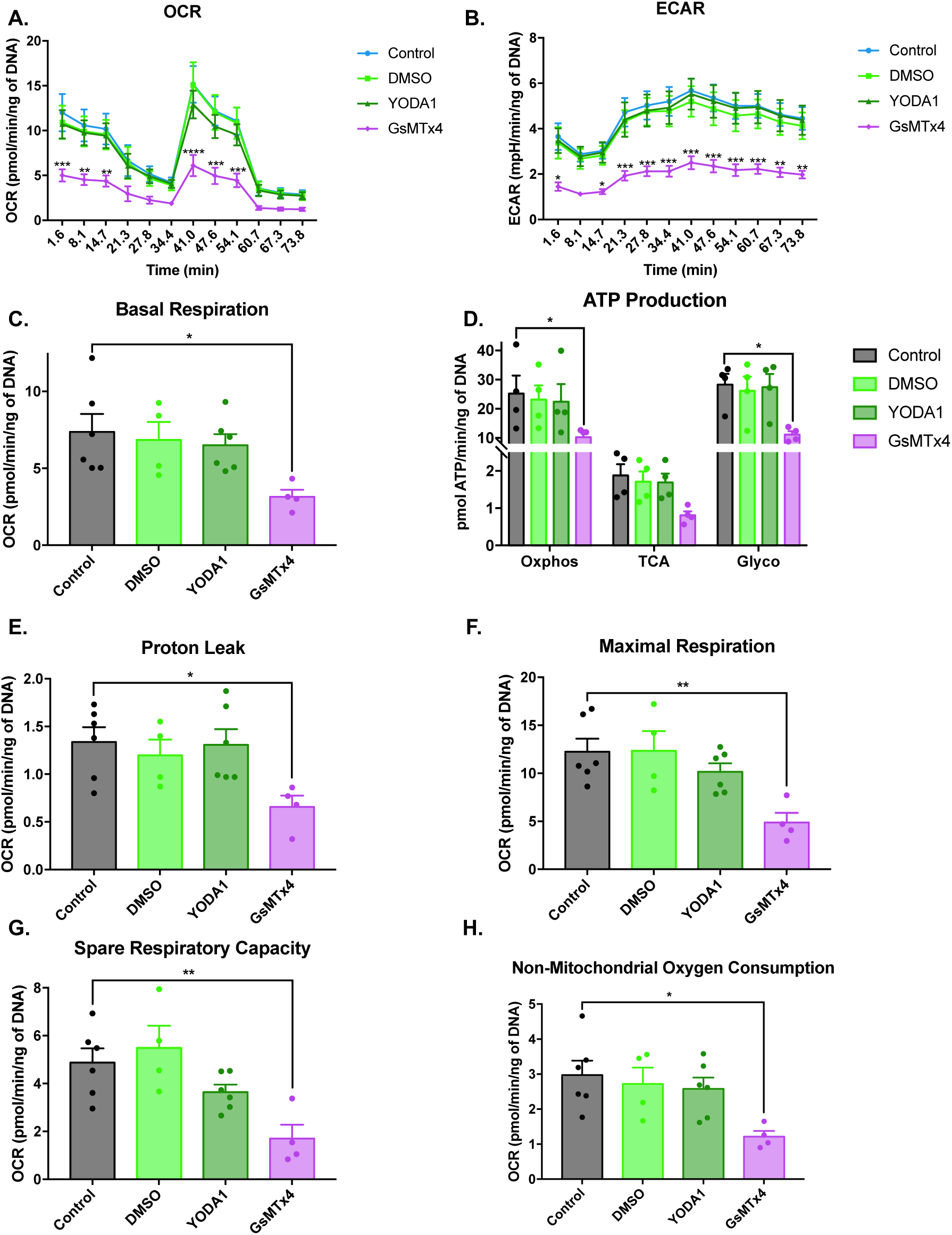
PIEZO1 inhibition affects the oxygen consumption and extracellular acidification rate in VM populations through impairment of glycolysis processes. To understand the mechanism of action through which GsMTx4 and Yoda1 modulate astrocyte activation, an extracellular flux analysis investigating the mitochondrial health state was performed using the MitoStress Test. Analysis of the effect of PIEZO1 activation and inhibition on cellular oxygen consumption (OCR) **(A)** and extracellular acidification rate (ECAR) **(B)** relative to cells cultured under control conditions. Analysis of the effect of PIEZO1 activation and inhibition on basal respiration **(C)**, ATP production **(D)**, proton leak processes **(E)**, maximal respiration **(F)**, the spare respiratory capacity **(G)** and non-mitochondrial oxygen consumption **(H)** relative to VM populations cultured under DMSO alone or control conditions. Data are represented as mean ± SEM (n=4-6). Two-way ANOVA with Tukey post hoc test was performed. *, **, ***, **** represents a statically significant difference versus control, (p<0.05), (p<0.01), (p<0.001) and (p<0.0001), respectively.

Specifically, inhibition of PIEZO1 by GsMTx4 led to a significant ∼57% (p=0.0364) reduction in the mitochondria basal respiration relative to cells cultured under control conditions, while activation by Yoda1 did not modulate basal respiration **(Figure 8C)**. Remarkably, GsMTx4 exposure significantly reduced ATP productions in VM cells. In particular, the oxidative phosphorylation pathway (Oxphos), the Krebs cycle (TCA) and the glycolysis cycle (Glyco) were all significantly downregulated by ∼59% (p=0.0324), ∼56% (n.s) and ∼61% (p=0.011), respectively, relative to VM cells cultured under control conditions **(Figure 8D)**. Similarly, maximal respiration was significantly impaired by PIEZO1 inhibition inducing a significant ∼60% (p=0.0073) reduction relative to VM cells cultured under control conditions **(Figure 8F)**. The spare respiratory capacity was also significantly modulated by PIEZO1 inhibition with a decrease of ∼65% (p=0.0091) relative to VM cells cultured under control conditions **(Figure 8G)**. Interestingly, the non-mitochondrial oxygen consumption was also affected in VM populations exposed to the PIEZO1 antagonist, revealing a significant reduction of ∼59% (p=0.0227) relative to VM cells cultured under control conditions **(Figure 8H)**. PIEZO1 inhibition also induced a significant ∼50% (p=0.04) decrease in mitochondrial proton leak relative to VM cells cultured under control conditions **(Figure 8E)**. However, it was noted that neither DNA content nor the mitochondria bioenergetic health index was significantly modulated in any of the control or experimental groups, implying the principal reduction in mitochondria function was independent of cell loss or disruption to cellular health processes **(Figure S8D,E)**. Collectively, these findings suggest that the induction of a reactive astrocyte phenotype via GsMTx4 inhibition of PIEZO1 is accompanied by glycolysis impairment and diminished ATP production.

## 4. Discussion

In this work, we provide evidence for the role of milliscale shear stress in promoting reactive gliosis, and the evolution of the fluid-filled space at the peri-electrode interface and propose a mechanism of action mediated by the mechanical activation of transmembrane ion channel PIEZO1 **(Figure 9)**. Specifically, we developed an *in silico* model and show that respiration induced electrode micromotions are translated into milliscale shear stress and propagated through the peri-electrode fluid-filled space to the receding neural tissues **(Figure 1)**. Critically, we show that the shear stress amplitude is inversely proportional to the electrode-tissue distance and suggest that the peri-electrode fluid space reaches a steady state when a tissue shear stresses of <0.1 Pa is achieved through tissue regression.

**Figure 9.**
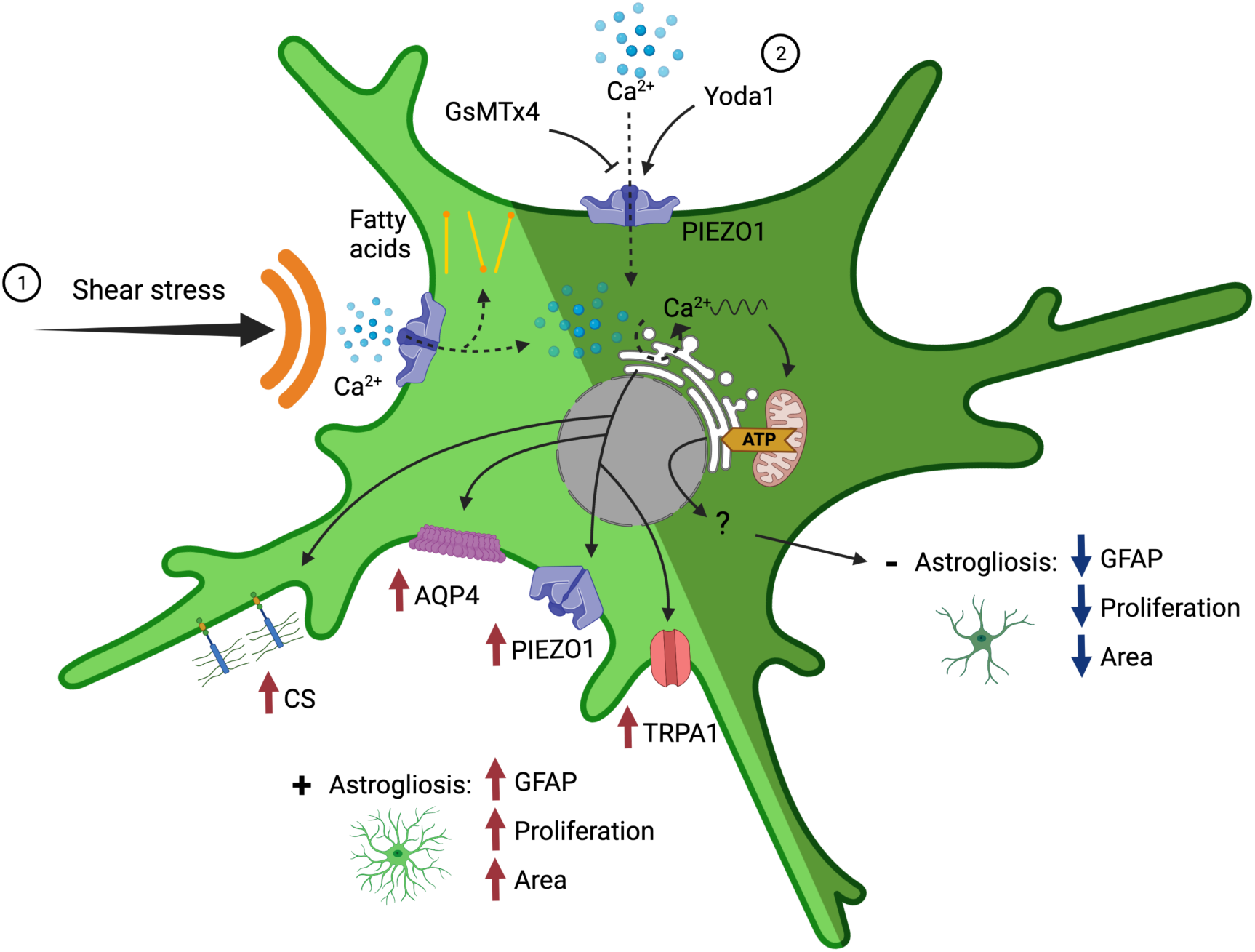
Proposed mechanisms of shear stress-mediated astrocyte activation and PIEZO1 inhibition with GsMTx4 or its overactivation with Yoda1 leading to the exacerbation or rescue of astrogliosis. **(1)** Shear stress applied to astrocytes, either through in vitro oscillatory fluid flow stimulation or in vivo electrode micromotion, opens the PIEZO1 mechanosensitive ion channel which allows calcium ions to flow inside the cells and initiating intracellular calcium waves triggering the overexpression of the glial fibrillary acid protein (GFAP), chondroitin sulfates (CS), aquaporin-4 (AQP4), Piezo type mechanosensitive ion channel component 1 (PIEZO1), transient receptor ankyrin 1 (TRPA1) and leading to an increased cell area and proliferation. **(2)** Overactivation of the PIEZO1 channel using the agonist Yoda1 results in an increased Ca^2+^ influx initiating ER-stored calcium waves which induces a rise in mitochondria mediated glycolysis and increased ATP production which results in a reduction of the astrogliosis process. Schematic created with BioRender.com.

We went on develop an *in silico* informed *in vitro* model of neuroelectrode micromotion induced fluid shear stress derived from a parallel-plate flow chamber. We subsequently investigated the effects of physiologically relevant shear stress on multiple indicators of astrocyte reactivity, inflammation and neurogenesis **(Figure 2-4)**. Critically, we identified that PIEZO1 and TRPA1 became significantly upregulated *in vitro* in response to fluid shear stress and went on to confirm these upregulations in an *in vivo* model of gliosis **(Figure 5-6)**. Finally, we demonstrated that chemical activation of PIEZO1 attenuated gliosis and stimulated neuronal regeneration *in vitro* Conversely, inhibition of PIEZO1 promoted a reactive astrocyte phenotype **(Figure 7)**. Finally, we showed that PIEZO1 inhibition impaired mitochondrial functions including ATP production, respiration, and glycolysis **(Figure 8)**.

Previous *in vivo* studies have demonstrated conclusively that micromotion induced peri-electrode shear stress contributes significantly to the development of glial scarring [33–36]. Furthermore, a dynamic process of tissue recession from the electrode and a reduction in device integration is frequently observed at the peri-electrode region weeks to months following implantation [1]. This tissue recession leads to the development of a growing peri-electrode interstitial fluid-filled space representing the *de novo* neuroelectrode interface which reaches up to 4 times the diameter of the implanted device [33,34,110] and is filled by an interstitial fluid composed by an inhomogeneous mix of cerebrospinal fluid and debris of cells and extracellular matrix, with an average viscosity of ∼30 mPa.s^−1^ [87]. Furthermore, this growing extracellular space has been shown to be exacerbated in devices fixed to the skull [33,34] due an increase in relative micromotion [35,45].

It can be hypothesised that the fluid within the peri-electrode void undergoes localised flow in response to electrode micromotion, imparting shear stress onto the peri-electrode tissues, leading to cell death and void expansion before reaching a steady state electrode-tissue distance, at which shear stresses no longer induce necrosis but chronic astrocyte reactivity, leading to glial scar development. Critically, we show that the evolution of this space is independent of electrical stimulation.

The underlying molecular mechanisms however of peri-implant gliosis remains unclear [1] and research has failed to develop comprehensive *in vitro* models which recapitulate the complex glial scarring process [151]. Most current *in vitro* models of gliosis induce glial cell proliferation and activation, as well as trigger neurodegeneration, however they do not elicit concurrent or chronic markers of glial scar development [83–85,151]. In this study, a physiologically representative oscillatory fluid flow shear stress model was found to trigger astrogliosis and neuronal loss, coupled with an overexpression of CS and a downregulation of the glypican 5 gene. This is relevant as *in vivo* studies of glial scar tissue have revealed a disbalance of heparan/chondroitin sulfates distribution, which inhibit neuronal growth and axonal regeneration as well as perpetuate inflammation [7,152–154]. Furthermore, the fluid shear stress model induced an overexpression of the water channel aquaporin-4 (AQP4) in VM astrocytes, an established marker of astrocyte reactivity and initiator of fibrosis through perturbation of transcellular water transport mechanisms [112,155– 157].

The effects of *in vitro* shear stress on the pro-inflammatory state of VM cell populations was further validated via a protein microarray and modulation to GFAP and ß-tubulin III expression, as well as decreased Myelin synthesis, indicating a loss of neuron and axon integrity were noted [158–160]. Moreover, proteomic analysis also revealed increases in the synthesis of establish brain injury/inflammation markers, including Nestin [161–163], Olig2 [164–166], CD81 [167,168] and OX42 [169,170].

*In vitro* fluid flow shear stress was also shown to initiate genomic changes resembling those reported in brain injury and neuroinflammatory contexts. Specifically, the upregulation of Casp3 and App, coupled with a simultaneous decrease in the expression of ACTB, depicts the onset of the neuronal apoptosis process [113–117,144], which was further indicated by the downregulation of neurogenesis-related genes Neurod1, Robo1, Ywhaq and HMCN1 [132,133,142,143,171–173]. Furthermore, the increased expression of two astrocytosis specific genes, S100a6 and S100B, points to a proliferative state of the astroglial population [118–121], further asserted by the positive regulation of markers of cell development, differentiation, proliferation and cycle regulation CCN1, CCN2, CSNK1E, ACTG1, EFNB1, CRB2 and CRB3, which are also reported as associated with glioblastoma, brain injury and astrocyte reactivity [122,141,174].

Interestingly, VM cultures exposed flow shear stress conditions also exhibited upregulated expression of NRG1, PAFAH1B1, DCX and GDNF and downregulation of NDN, which could suggest a protective and/or survival response triggered by the neuronal population in reaction to increased proliferation of astrocytes [129–131,134,135,140,175–178]. Conversely, the upregulation of BMP4 has been shown to act as a suppressor of neurogenesis and to play a role in sustaining astrogenesis [136,137,179,180], which suggest a retro-control of this neuronal-rescue process. Interestingly, HIPK2 also exhibited downregulated expression which has been shown to occur with noxious stimuli (such as shear stress), to alter p53 expression and lead to neuron dysfunction [181]. In addition, the decreased expression of NPHP4 and PTPN14, suggests inhibition of the Yap/Taz complex and an activation of the hippo signalling pathway [145–147].

Interestingly, it is thought that the *in vitro* model did not contain microglia or oligodendrocytes (due to low numbers in the midbrain at this developmental stage [182] & [183]), suggesting that the neurodegeneration observed from 7 day post-stimulation was derived from an increasing astrocyte reactivity state. In fact, Liddelow & Barres proposed a mechanism where astrocytes, after injury, release neurotoxic factors [184], which have recently been described as saturated lipids [185]. This hypothesis correlates with our genomic data analysis which also predicted the release of fatty acids from VM populations following exposure to shear stress *in vitro* **(Figure S7B)**.

Mechanically gated ion channels have recently been implicated as playing key roles in the homeostasis of the CNS and in mediating mechanotransduction in neural populations [49,50,52,60,61]. Indeed several ion channels have been shown to play a role in the reception of mechanical cues and have been shown to become perturbed in the progression of multiple disease states [25,48,68]. In particular astrocytic PIEZO1 has been shown to be overexpressed in tissues with elevated beta-amyloid deposition and in glioma masses due to increased tissue rigidity [69,70,80]. TRPA1 has also been described as upregulated in astrocytes and Schwann cells following neural inflammation and injury [59,186]. Moreover, both TRPA1 and PIEZO1 have been implicated as mediators of the astrocyte neuroinflammatory response by modulating the synthesis of pro-inflammatory cytokines [64] or hormones [65].

Here, PIEZO1 overexpression in reactive astrocyte populations (as confirmed by elevated GFAP synthesis) was observed *in vitro* and at the peri-electrode site *in vivo*, indicating that these structures may play a role in mechanosensation of fluid-flow generated shear stress. It can further be hypothesised that PIEZO1 overexpression in astrocyte populations induces a cellular shift towards a pro-inflammatory phenotype, as demonstrated in macrophages by Atcha, et al. [187] and can play a role in establishing chronic peri-electrode inflammation [48].

Interestingly, in both *in vitro* and *in vivo* models, TRPA1 displayed a more complex expression profile; indeed, the TRPA1 receptor was upregulated in VM populations exposed to shear stress conditions, however, its presence was more abundant in astrocytes populations with a moderate GFAP content and spatially more distant from the vicinity of the electrode. This expression profile may due to the role of astrocytic TRPA1 in calcium induced glutamate release which has been shown to activate neighbouring neurons [148,149,188,189]. It can be hypothesised that TRPA1 expression is localised to those astrocyte populations which interact with neuronal cells and which are less abundant at the peri-electrode interface.

To better understand the role of MS ion channels, in both physiological and pathological contexts and to assess their potential as therapeutic targets, chemical inhibitor, GsMTx4, and activator Yoda1 can be employed to modulated PIEZO1 channel activation in vitro [51,54,64,69,190–193]. Here it was observed that GsMTx4-induced PIEZO1 inhibition led to astrocyte reactivity and neurodegeneration and potentiated the proinflammatory effects of fluid flow shear stress *in vitro.* Pathak and colleagues showed comparable results in neural stem cells, and both chemical or siRNA-mediated PIEZO1 inhibition stimulated astrogenesis and reduced neurogenesis through a decrease in Yap mediated nucleo-translocation [51]. Moreover, H. Liu et al. revealed that the inhibition of PIEZO1 increases the release of pro-inflammatory mediators in microglia [194], again suggesting a pro-gliosis effect of PIEZO1 inhibition.

Conversely, PIEZO1 activation through Yoda1 supplementation, was observed to negate the pro-inflammatory effects triggered by oscillatory fluid shear stress. This is in agreement with a previous study by Velasco-Estevez et al., who showed that the activation of PIEZO1 in LPS exposed cortical astrocytes led to a reduction in the synthesis of pro-inflammatory cytokines and reduced gliosis *in vitro* [64]. Several further studies have also showed that PIEZO1 is upregulated after axon/neuronal injury and that its loss or inhibition boost axonogenesis and regeneration [54,195– 197], suggesting multiple and contradictory roles for the same mechanosensitive ion channel PIEZO1 according to the cell type and to physiological context.

This duality was also observed in this study, where fluid shear stress induced an upregulation of PIEZO1 expression, simultaneously to astrocyte activation and neural degeneration *in vitro*, effects which were reversed by chemical activation of PIEZO1. Interestingly, a recent study by Swain, et al. has shown that PIEZO1 activation through either shear stress or Yoda1, leads to increased synthesis of PLA2 in pancreatic cells, an enzyme involved in fatty acid release (an effect predicted by our genomic data), which triggers the opening of the TRPV4 channel and perpetrates dysregulated calcium flux, resulting in mitochondria dysfunction [198]. Subsequently, a growing body of research has shown that restoration of astrocytes and neural cell metabolism following CNS traumatic injury or disease, can produce a neuroprotective effect and reduce coinciding astrogliosis [199–202].

Recently, several studies have linked mechanosensing [203], MS ion channels [204] and particularly the activation of PIEZO1 with an increased release of ATP [190,205– 209]. To investigate the role of PIEZO1 in regulating neural metabolism *in vitro*, we combined metabolic flux analysis with PIEZO1 chemical modulation and observed that, GsMTx4-induced PIEZO1 inhibition increased pro-inflammatory process and decreased critical mitochondrial functions in VM cell populations, including glycolysis and ATP production, effects which were not observed with Yoda1 mediated PIEZO1 activation. Interestingly, Lopez-Fabuel et al. showed that decreased mitochondrial function and increased ROS production, promotes astrocyte survival while negatively influencing neuron viability [210]. This effect was further described by Weber and Barros, who discussed the “selfishness” of reactive astrocytes, observing that following injury or in response to disease, activated astrocytes will alter their metabolism and reduced their neuronal support functions to favour their own survival [211,212].

Together, these data indicate that PIEZO1 activation impacts mitochondrial functions and cellular metabolism, which are crucial for the homeostasis of neural tissues [213,214] and play an important role in controlling both neuron and astrocyte functions [51]. Mechanistically, it can be hypothesised that shear stress-induced PIEZO1 activation, promotes a sustained augmentation of intracellular calcium transportation processes which leads to the release of fatty acids and to the impairment of mitochondrial functions, drastically reducing ATP production and promoting neuronal death and astrocyte reactivity **(Figure 9)**.

In conclusion, we have shown that neuroelectrode micromotion is translated into milliscale shear stress at the tissue-electrode interface and hypothesise that this mechanism gives rise to the peri-electrode fluid filled space. Our work suggests that the mechanosensitive ion channels PIEZO1 and TRPA1 are key mediators of astrogliosis and act as regulators of critical metabolic process in neural populations. Finally, it may be inferred that electrode functionalisation with chemical agonist/antagonist of Piezo1 may promote chronic electrode stability in vivo.

## 5. Acknowledgments

This work has been funded through Science Foundation Ireland (SFI) and the European Regional Development Fund (grant number 13/RC/2073). The authors acknowledge the facilities and scientific and technical assistance of the Centre for Microscopy & Imaging at the University of Galway (www.imaging.nuigalway.ie). The authors would like also to thank Prof David Hoey from Trinity College Dublin (Ireland) for allowing the use of his parallel flow chamber design and Prof Abhay Pandit from the University of Galway for the financial support.

## 7. Authorship contribution statement

**Alexandre Trotier**: Conceptualisation, Writing – original draft, Data curation, Writing – review & editing, Formal analysis, Investigation, Methodology.

**Enrico Bagnoli**: Formal analysis, Data curation, Investigation, Methodology.

**Tomasz Walski**: Formal analysis, Data curation, Investigation, Methodology.

**Judith Evers**: Formal analysis, Data curation, Investigation, Methodology.

**Eugenia Pugliese**: Formal analysis, Data curation, Investigation, Methodology.

**Madeleine Lowry**: Resources, Writing – review & editing.

**Michelle Kilcoyne**: Resources, Writing – review & editing.

**Una Fitzgerald**: Supervision, Resources, Writing – review & editing.

**Manus Biggs**: Supervision, Funding acquisition, Resources, Project administration, Conceptualised, Writing – original draft, Writing – review & editing.

## Abbreviations

ADORA1: Adenosine A1 Receptor
ACTB: actin beta
ACTG1: actin gamma 1
AMOT: angiomotin
AMOTL2: angiomotin like 2
APP: amyloid beta protein precursor
APOE: apolipoprotein E
AQP4: aquaporin 4
ATP: adenosine triphosphate
BCL2: BCL2 Apoptosis Regulator
BDNF: Brain Derived Neurotrophic Factor
BHI: bioenergetic health index
BMP4: Bone Morphogenetic Protein 4
Casp3: caspase 3
CCNE1: cyclin E1
CCNE1: cyclin E2
CFD: computational fluid dynamics
CNS: central nervous system
CRB2: Crumbs Cell Polarity Complex Component 2
CRB3: Crumbs Cell Polarity Complex Component 3
CS: chondroitin sulfates
CSNK1D: Casein Kinase 1 Delta
CSNK1E: Casein Kinase 1 Epsilon
CXCL2: C-X-C Motif Chemokine Ligand 2
DAPI: 4’,6-diamidino-2-phenylindole
DBS: deep brain stimulation
DCX: doublecortin
DRD2: Dopamine Receptor D2
ECAR: extracellular acidification rate
EFNB1: Ephrin B1
EGF: Epidermal Growth Factor
ERBB2: Erb-B2 Receptor Tyrosine Kinase 2
FGF2: Fibroblast Growth Factor 2
FJX1: Four-Jointed Box Kinase 1
FVM: finite volume model
GDNF: Glial Cell Derived Neurotrophic Factor
GFAP: glial fibrillary acidic protein
GPC5: glypican 5
HIPK2: Homeodomain Interacting Protein Kinase 2
HMCN1: hemicentin 1
HPRT1: Hypoxanthine Phosphoribosyltransferase 1
Iba1: Ionised calcium-binding adaptor 1
MS: mechanosensitive
MYC: Fibroblast Growth Factor 2
NDN: necdin
NeuN: Neuronal nuclear antigen
Neurod1: Neurogenic differentiation factor 1
NOG: noggin
NPHP4: Nephrocystin 4
NRG1: Neuregulin 1
OCR: oxygen consumption rate
Olig2: Oligodendrocyte transcription factor
OxPhos: Oxidative phosphorylation
PAFAH1B1: Platelet Activating Factor Acetylhydrolase 1b Regulatory Subunit 1
PIEZO1: Piezo-type mechanosensitive ion channel component 1
PIEZO2: Piezo-type mechanosensitive ion channel component 2
PLL: poly-L-lysine
PPFC: parallel plate flow chamber
PPP2CA: Protein Phosphatase 2 Catalytic Subunit Alpha
PTN: pleiotrophin
PTPN14: Protein Tyrosine Phosphatase Non-Receptor Type 14
ROBO1: Roundabout Guidance Receptor 1
ROS: reactive oxygen species
SHH: Sonic Hedgehog Signalling Molecule
SOD1: Superoxide Dismutase 1
S100A6: S100 Calcium Binding Protein A6
S100B: S100 Calcium Binding Protein B
TCA: Tricarboxylic acid
TGFB1: Transforming Growth Factor Beta 1
TRAAK: TWIK-related Arachidonic Acid-Stimulated K+
TREK-1: Potassium channel subfamily K member 2
TRP: transient receptor potential cation channel
TRPA1: transient receptor potential ankyrin 1
TRPV1: transient receptor potential vanilloid receptor 1
VM: ventral mesencephalic
WSS: wall shear stress
YWHAQ: Tyrosine 3-Monooxygenase/Tryptophan 5-Monooxygenase Activation Protein Theta

