## Supplementary file for "Micromotion derived fluid shear stress mediates peri-electrode gliosis through mechanosensitive ion channels"

### Supplementary data

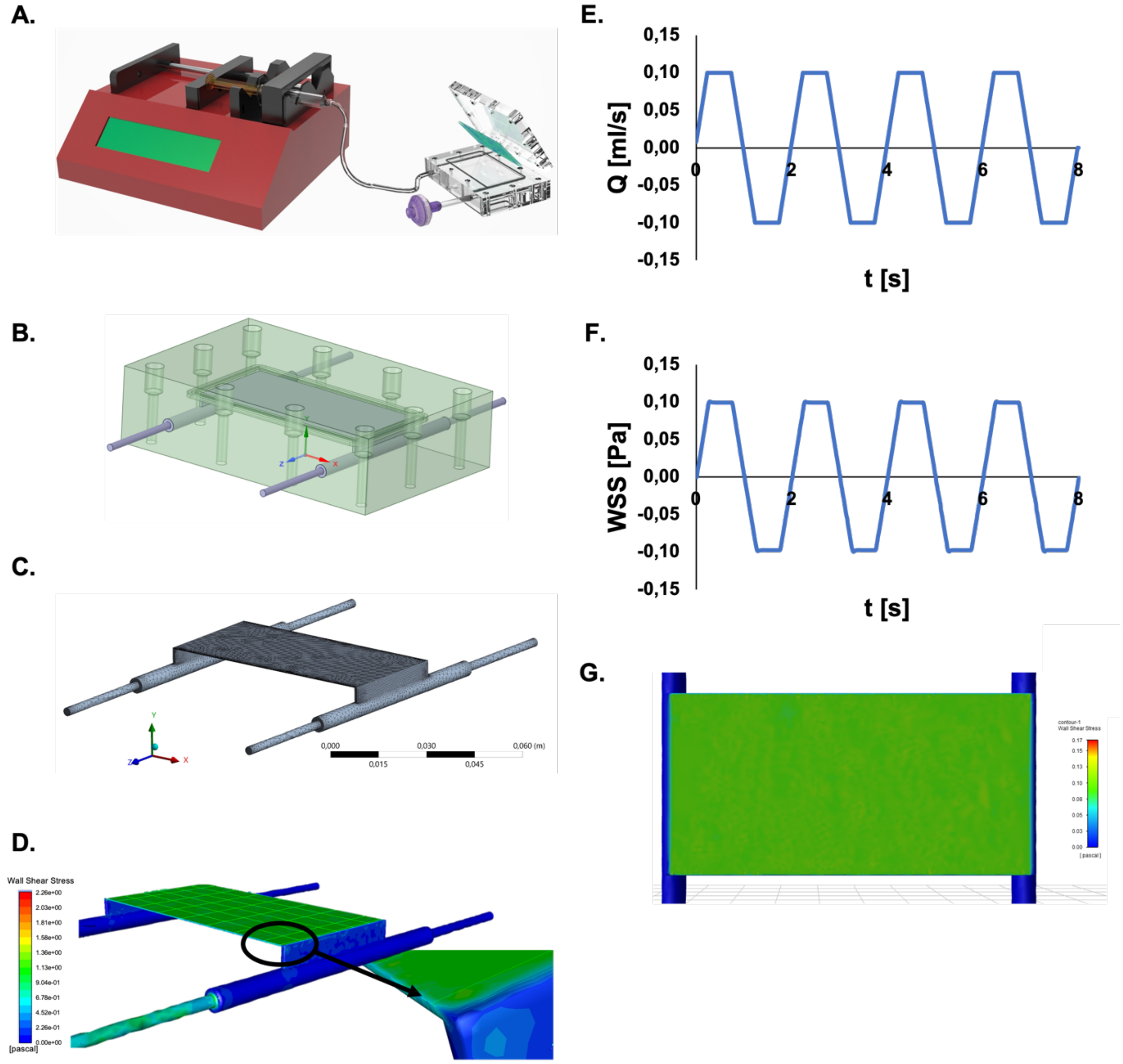


***Figure S1 Parallel plate flow chambers can induce homogeneous and consistent oscillatory fluid flow shear stresses on neural cell monolayers.*** *General concept of the setup providing oscillatory fluid stimulation conditions to VM cultures including PPFCs enclosing a glass slide covered with a cell monolayer connected to a syringe-pump which controls the flow direction and rate* ***(A)****. The 3D design of the PPFC geometry, where green represents solid parts and violet denotes fluid domain* ***(B)****. The mesh of the fluid volume used for numerical calculations* ***(C)****. WSS distribution on the surface of the PPFC with sampling lines used for the assessment of the coefficient of variance* ***(D)****. Changes in oscillatory volumetric flow rate in time at the inlet of the PPFC* ***(E)*** *with accompanying changes of WSS measured in the central point of the microchannel surface* ***(F)*** *for Q = 4.49 ml/min. The homogeneity of WSS on the top surface of the microchannel for Q = 4.49 ml/min during the steady flow period* ***(G)****.*

*Table S1 Summary of test conditions and results of computational studies of the WSS homogeneity in the developed PPFC system.*

| **Q [ml/min]** | 4,49 | 44,89 | 380,46 |
|---|---|---|---|
| **WSS [Pa] mean** | 0.098 | 0.990 | 9.70 |
| **WSS [Pa] SD** | 0.008 | 0.079 | 0.83 |
| **CV** | 8.4% | 8.0% | 8.6% |
| **p-value*** | < 0.0001 | < 0.0001 | < 0.0001 |

* results of the two-tailed unpaired t-test comparison of the WSS values at the top surface of the microchannel (full) vs. the surface reduced by the edges area corresponding to the decreased values of the WSS (reduced).

*Table S2: Antibody source and dilution used for immunohistochemistry*

| Antibody | Host/Isotype | Supplier | Catalogue No. | Dilution |
|---|---|---|---|---|
| **Primary** | | | | |
| GFAP | Mouse mAb | Sigma | G3893 | 1:500 |
| β-Tubulin III | Rabbit pAb | Sigma | T2200 | 1:500 |
| Chondroitin Sulfate | Mouse mAb | Sigma | C8035 | 1:200 |
| AQP4 | Rabbit pAb | Alomone | AQP-004 | 1:500 |
| PIEZO1 | Rabbit pAb | Alomone | APC-087 | 1:200 |
| TRPA1 | Rabbit pAb | Alomone | ACC-037 | 1:200 |
| NeuN | Rabbit mAb | Abcam | ab177487 | 1:1000 |
| Iba1 | Rabbit pAb | Wako | 019-19741 | 1:500 |
| **Secondary (Alexafluor)** | | | | |
| Anti-mouse 488 | Donkey pAb | Invitrogen | A-21202 | 1:500 |
| Anti-Rabbit 594 | Donkey pAb | Invitrogen | A-21207 | 1:500 |

*Table S3:Summary of all commercial antibodies used for the construction of the antibody microarray*

| Order | Probe | Concentration (mg/mL) | Species | Company | Ref |
|---|---|---|---|---|---|
| 1 | Integrin beta5 | 0,35 | Rabbit | Cell signalling | D24A5 |
| 2 | P-Smad1 | 0,25 | Rabbit | Cell signalling | S206 D40B7 |
| 3 | P-p44/42 MAPK | 0,05 | Rabbit | Cell signalling | T202/Y204 (197G2) |
| 4 | b-catenin 200 ug | 1 | Rabbit | Millipore | 06-734 |
| 5 | Anti-Active-b-Catenin (anti ABC) Antibody, clone 8E7 | 1 | Mouse | Sigma® | 05-665-25UG |
| 6 | Integrin beta1 [EPR1040Y] 100ul ((0.157 mg/ml) | 1 | Rabbit | Abcam | ab134179 |
| 7 | Collagen I [EPR7785] 100 ul (0.875 mg/ml) | 1,003 | Rabbit | Abcam | ab138492 |
| 8 | Anti-Mouse IgG H&L | 0,05 | Goat | Abcam | ab150113 |
| 9 | FAK 100ug | 0,547 | Mouse | MBL | 12G4 |
| 10 | BMPR1A (PA5-11856) | 1 | Rabbit | ThermoFisher | PA5-11856 |
| 11 | Collagen II [2B1.5] 250 ul (0.2 mg/ml) | 0,25 | Mouse | Abcam | ab185430 |
| 12 | Myelin [MBP101] 100 ug (3.2 mg/ml) | 0,125 | Mouse | Abcam | ab62631 |
| 13 | Collagen V 100 ug (1mg/ml) | 0,856 | Rabbit | Abcam | ab7046 |
| 14 | SCXA (100ug) | 0,1 | Rabbit | Abcam | ab58655 |
| 15 | Biglycan (100ug) | 0,183 | Rabbit | Abcam | ab49701 |
| 16 | TBHS4 | 1,086 | Rabbit | Abcam | ab176116 |
| 17 | Tenascin C (50ug) | 0,1 | Rabbit | Abcam | ab88280 |
| 18 | Decorin | 1,075 | Rabbit | Abcam | ab175404 |
| 19 | Tenomodulin (100ug) | 0,5 | Rabbit | Abcam | ab203676 |
| 20 | Collagen III 100 ug ( 1mg/ml) | 0,002 | Rabbit | Abcam | ab7778 |
| 21 | Olig2 [EPR2673] | 0,25 | Rabbit | Abcam | ab109186 |
| 22 | KCNK4 100 ul (0.5mg/ml) | 0,002 | Rabbit | Abcam | ab81367 |
| 23 | Integrin beta 3 [crc54] | 0,1 | Rabbit | Abcam | ab34409 |
| 24 | P-FAK | 0,002 | Rabbit | Cell signalling | Y925 |
| 25 | p-smad1/5 | 0,1 | Rabbit | Cell signalling | S463/465 |
| 26 | p44/42 MAPK (ERK 1/2) | 0,1 | Mouse | Cell signalling | L34F12 |
| 27 | Smad1 | 0,05 | Rabbit | Cell signalling | D59D7 |
| 28 | Smad5 | 0,004 | Rabbit | Cell signalling | D4G2 |
| 29 | Anti-beta Actin antibody | 0,75 | Mouse | Abcam | ab8226 |
| 30 | Anti-Glial Fibrillary Acidic Protein (GFAP) | 0,25 | Mouse | Sigma® | G3893 |
| 31 | Nestin (Rat-401) | 0,5 | Mouse | SantaCruz® | sc-33677 |
| 32 | CD81 (H-121) | 1 | Rabbit | SantaCruz® | sc-9158 |
| 33 | Integrin alphaM (OX42) | 1 | Mouse | SantaCruz® | sc-53086 |

| | | | | | |
|---|---|---|---|---|---|
| 34 | Phosphate buffered saline, pH 7.4 | | | | |
| 35 | Paxillin antibody [Y113] | 0,5 | Rabbit | Abcam | ab32084 |
| 36 | cleaved spectrin alpha II (h1186) | 0,5 | Rabbit | SantaCruz® | sc-23464 |
| 37 | Anti-Smad3 antibody [EP568Y] | 1 | Rabbit | Abcam | ab40854 |
| 38 | Anti-Chondroitin Sulfate antibody | 0,1 | Mouse | Sigma® | C8035 |
| 39 | Phosphate buffered saline, pH 7.4 | | | | |
| 40 | VR1 (H-150) | 1 | Rabbit | SantaCruz® | sc-20813 |
| 41 | Anticorps L-type Ca++ CP α1C (H-280) | 0,5 | Rabbit | SantaCruz® | sc-25686 |
| 42 | PIEZO1 Antibody (N-15) | 1 | Goat | SantaCruz® | sc-164319 |
| 43 | PIEZO2 Antibody (G-20) | 1 | Rabbit | SantaCruz® | sc-84763 |
| 44 | ANKTM1 (C-19) | 0,5 | Goat | SantaCruz® | sc-32353 |
| 45 | TREK-1 Antibody (C-20) | 1 | Goat | SantaCruz® | sc-11557 |
| 46 | Anti-β-Tubulin III | 0,75 | Rabbit | Sigma® | T2200 |

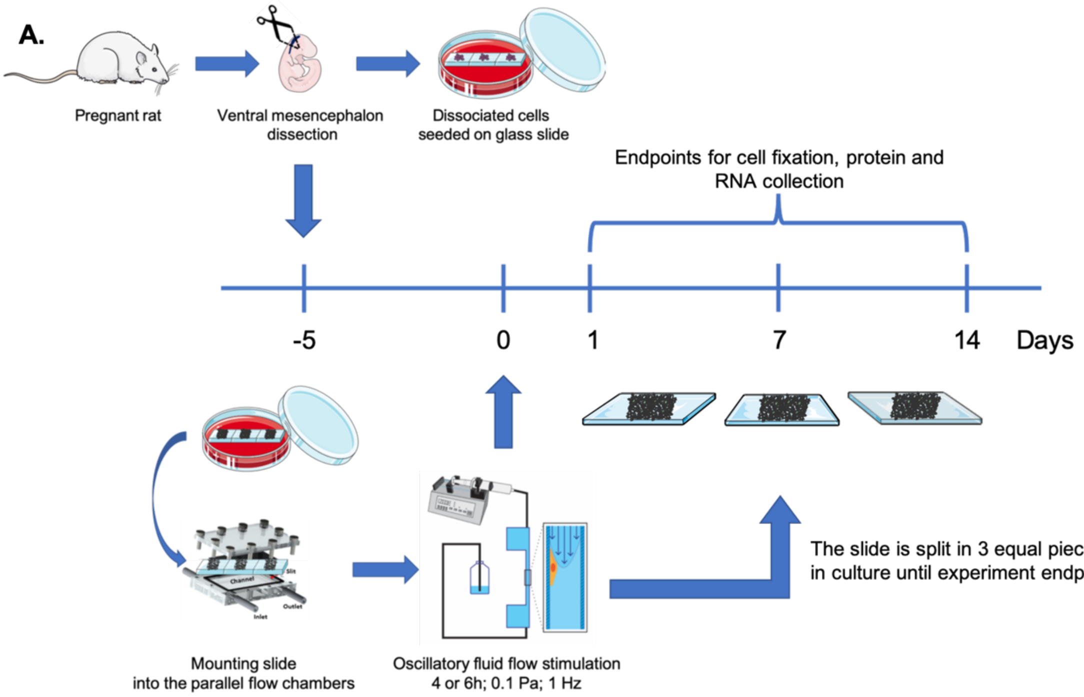


***Figure S2 Study timeline and setup.*** *Primary neural cells were extracted from the ventral mesencephalon of E14 rat embryos and seeded onto a glass slide five days prior to exposure to millipascal shear stress. At day 0, the glass slide was placed in the parallel flow apparatus and exposed to an oscillatory fluid flow at 0.1 Pa, 0.5 Hz for either 4 or 6h using pulsed culture medium. Following shear stimulation, glass slides were subdivided into 3 equal pieces, and kept in culture until experiment endpoint for either fixation or RNA/protein extraction.*

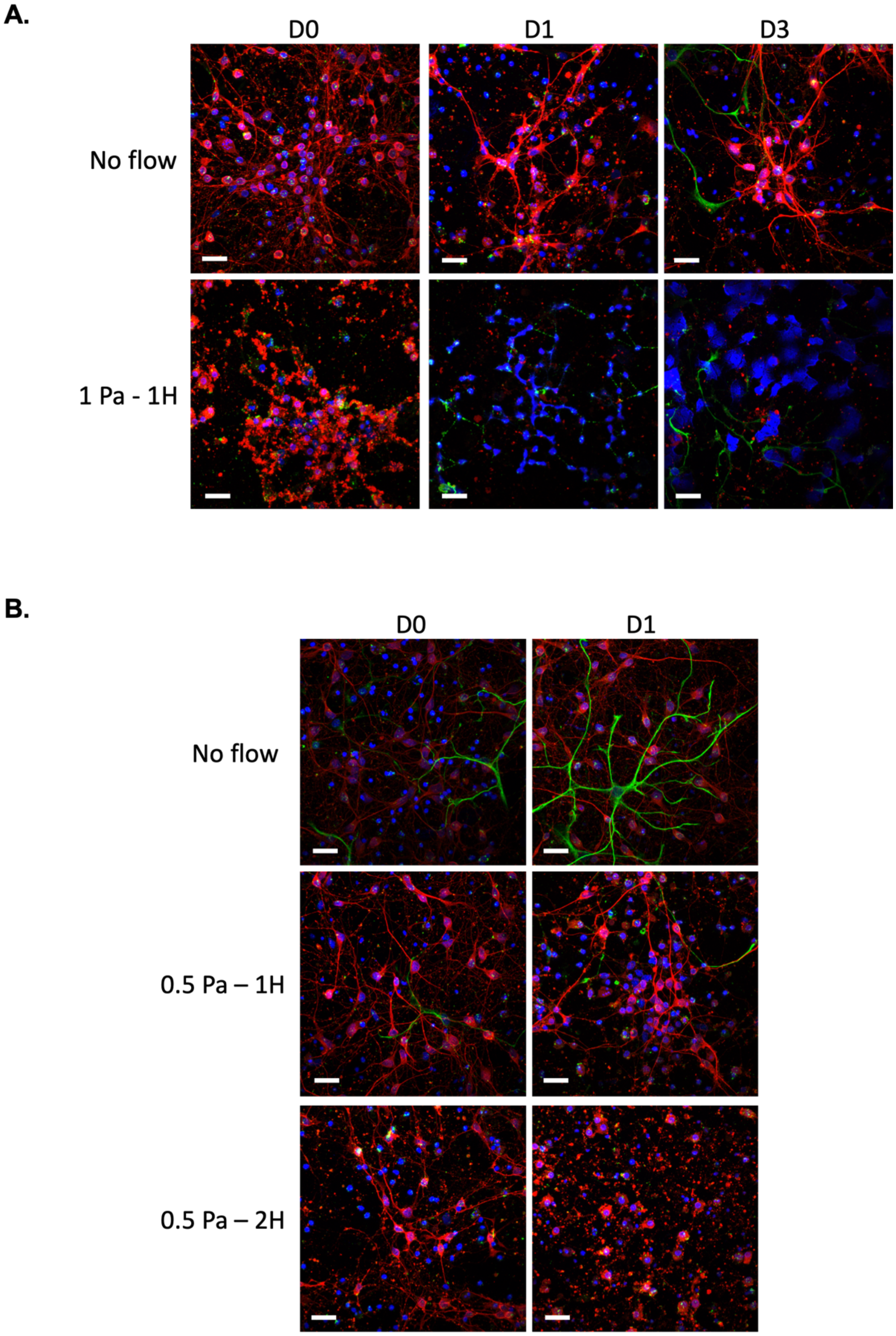


***Figure S3 The effect of oscillatory fluid flow shear stress magnitude (Pa) and duration on neuron viability.*** *Ventral mesencephalic cells could not survive a shear stress equals to 1 Pa for 1h* ***(A)*** *or 0.5 Pa during 2h* ***(B)****. Indeed, at day 0 immediately following the stimulation of 1Pa for 1h, the cytoplasm of all the cells appears to have burst and the culture did not recover over the 3 next days in culture* ***(A)****. Similarly, after a fluid flow stimulation of 0.5Pa during 2h, the VM culture seems severely deteriorated at day 0 in comparison to the static control and this deterioration seems exacerbated and to have led to complete loss of the cells at day 1 post-stimulation* ***(B)****. Scale bar = 20 µm.*

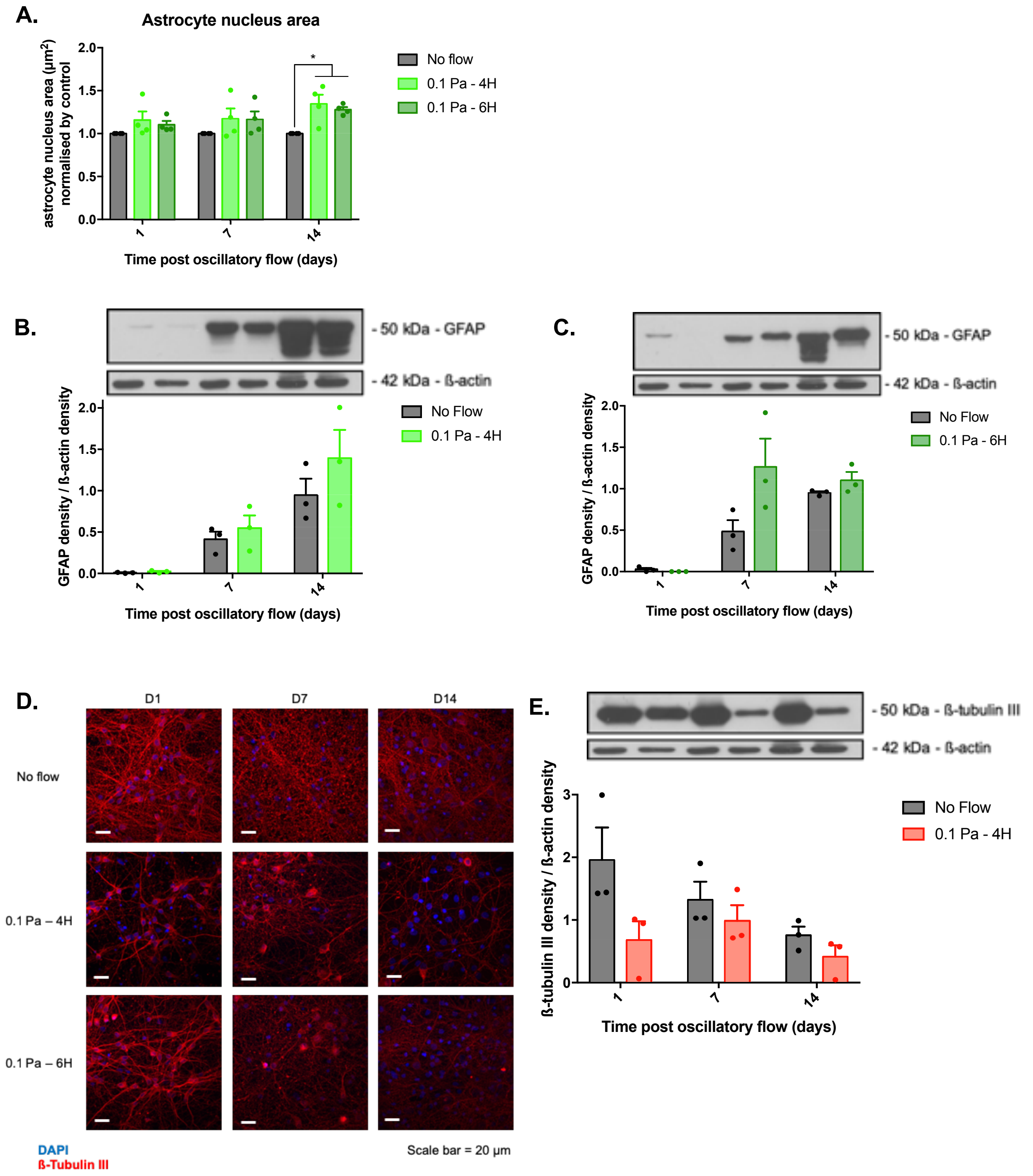


***Figure S4 Oscillatory fluid flow shear stress leads to astrogliosis and neurodegeneration in ventral mesencephalic (VM) cells.*** *The astrocyte nucleus average area also showed an increase of 15 to 25% at every time point in all stimulated conditions* ***(A)****. GFAP upregulation was confirmed by western blotting where the inducement of flow shear stress seems to induce an overexpression of the total GFAP protein after 7 and 14 days post stimulation* ***(B,C)****. VM cells' ß-tubulin III staining only* ***(D)*** *(scale bar = 20 µm; n=3). The shear stress effect on the neuronal cytoskeletal protein was confirmed by western blotting exhibiting a drop of the total protein expression in ß-tubulin III at every time points for the 4h stimulation at 0.1 Pa condition* ***(D)****. Data are represented as mean ± SEM (n=3-4). One-way ANOVA with Tukey post hoc test was performed. *, **, *** represents a statically significant difference (p<0.05), (p<0.01) and (p<0.001), respectively.*

**A.**

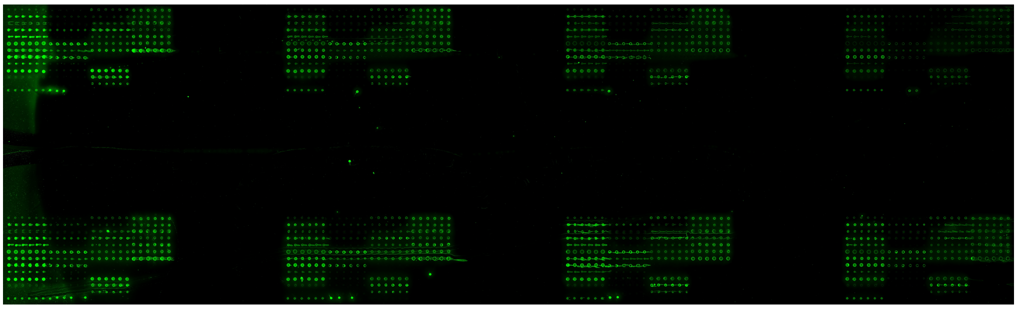


**B.**

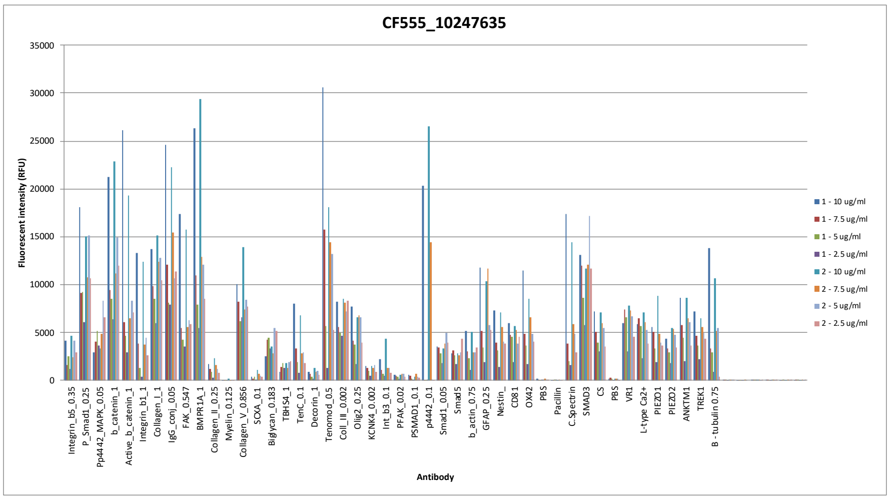


**C.**

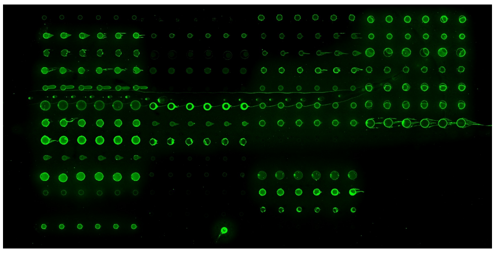

**D.**

| | | | |
|---|---|---|---|
| Integrin-ß-5 | SCXA | Smad1 | TRPV1 |
| P-Smad1 | Biglycan | Smad5 | L-type Ca++ |
| P-p44/42 MAPK | TBHS4 | ß-actin | PIEZO1 |
| ß-catenin | Tenascin | GFAP | PIEZO2 |
| Anti-Active-ß-Catenin | Decorin | Nestin | TRPA1 |
| Integrin ß-1 | Tenomodulin | CD81 | TREK-1 |
| Collagen I | Collagen III | OX42 | Anti-β-Tubulin III |
| Anti-Mouse IgG | Olig2 | PBS | |
| FAK | TRAAK | Paxillin | |
| BMPR1A | Integrin-ß-3 | cleaved spectrin α-II | |
| Collagen II | P-FAK | Anti-Smad3 | |
| Myelin | p-smad1/5 | Anti-Chondroitin Sulfate | |
| Collagen V | p44/42 MAPK | PBS | |

***Figure S5 Sample concentration optimisation of the antibody microarray.*** *Two representative protein samples of the in vitro study were labelled using CF™ 555 and incubated on a microarray slide at four different concentrations (10; 7.5; 5 and 2.5 µg/ml) and the image of the scanned slide* ***(A)*** *after incubation and washing was used to plot the fluorescence intensity of every printed antibody* ***(B),*** *in order to select the optimal protein concentration leading to the highest fluorescence reading without reaching saturation effects. A concentration of 7.5 µg/ml* ***(C)*** *was found optimal and sample 1 was systematically incubated as an internal control in the top right gasket of every slide of the study to ensure consistent printing and robust antibody performance. Each subarray were printed with antibodies in replicates of six spots as detailed in* ***(D)****.*

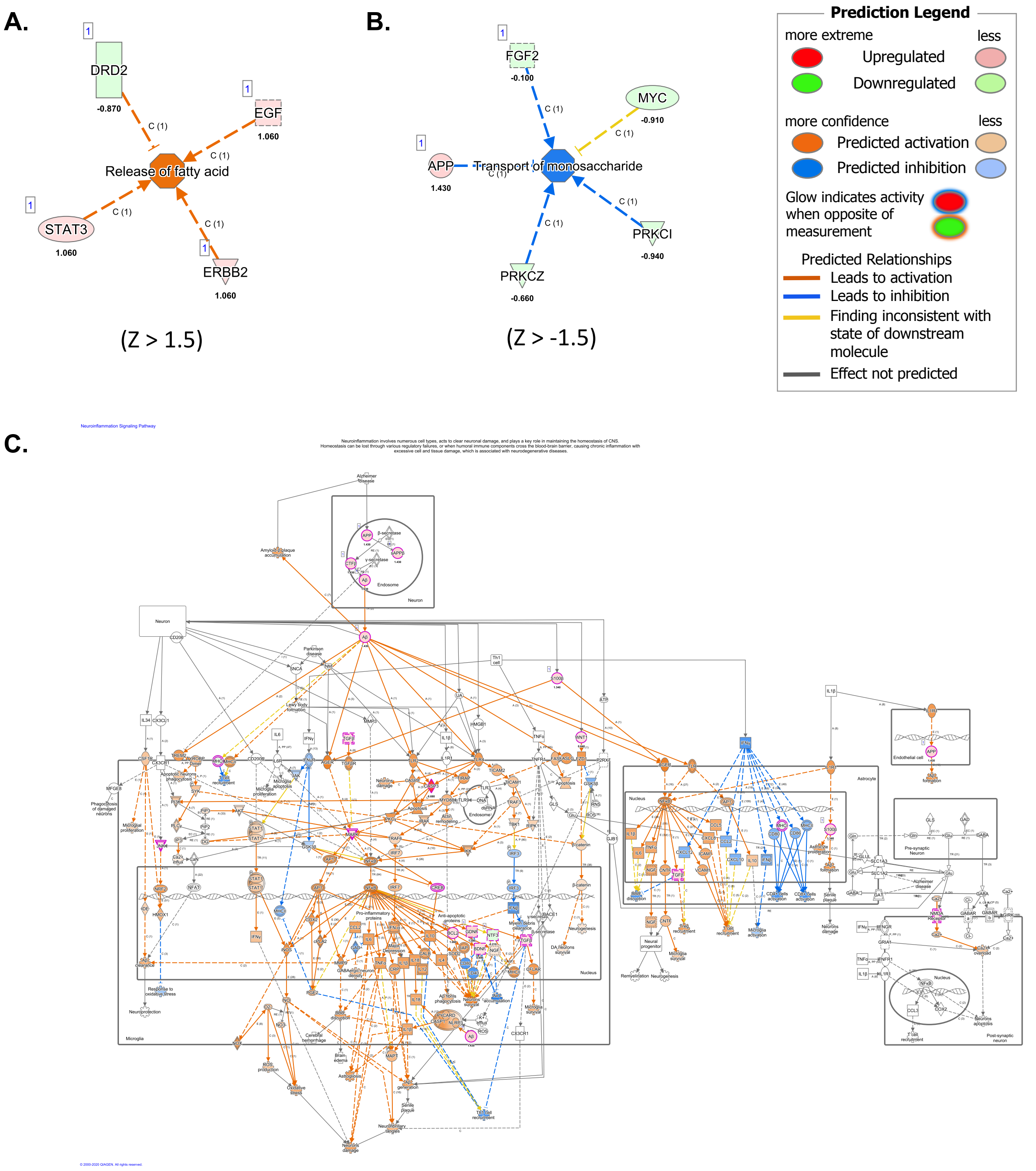


***Figure S6 Ingenuity Pathway Analysis predicted the activation of the neuroinflammation canonical pathway and biological functions related to metabolism***. *The Qiagen® IPA software predicted the activation of two additional biological function networks related to metabolism: the function "release of fatty acid" was predicted activated significantly modulated with an activation Z>1.5* ***(A)****, whereas the function "transport of monosaccharide" was detected inhibited with a Z< -1.5* ***(B)****. Moreover, the Qiagen® IPA software also predicted the overall activation of the neuroinflammation pathway, with a Z-score superior at 2, including 24 genes which underwent statistically significant modulation* ***(C)****.*

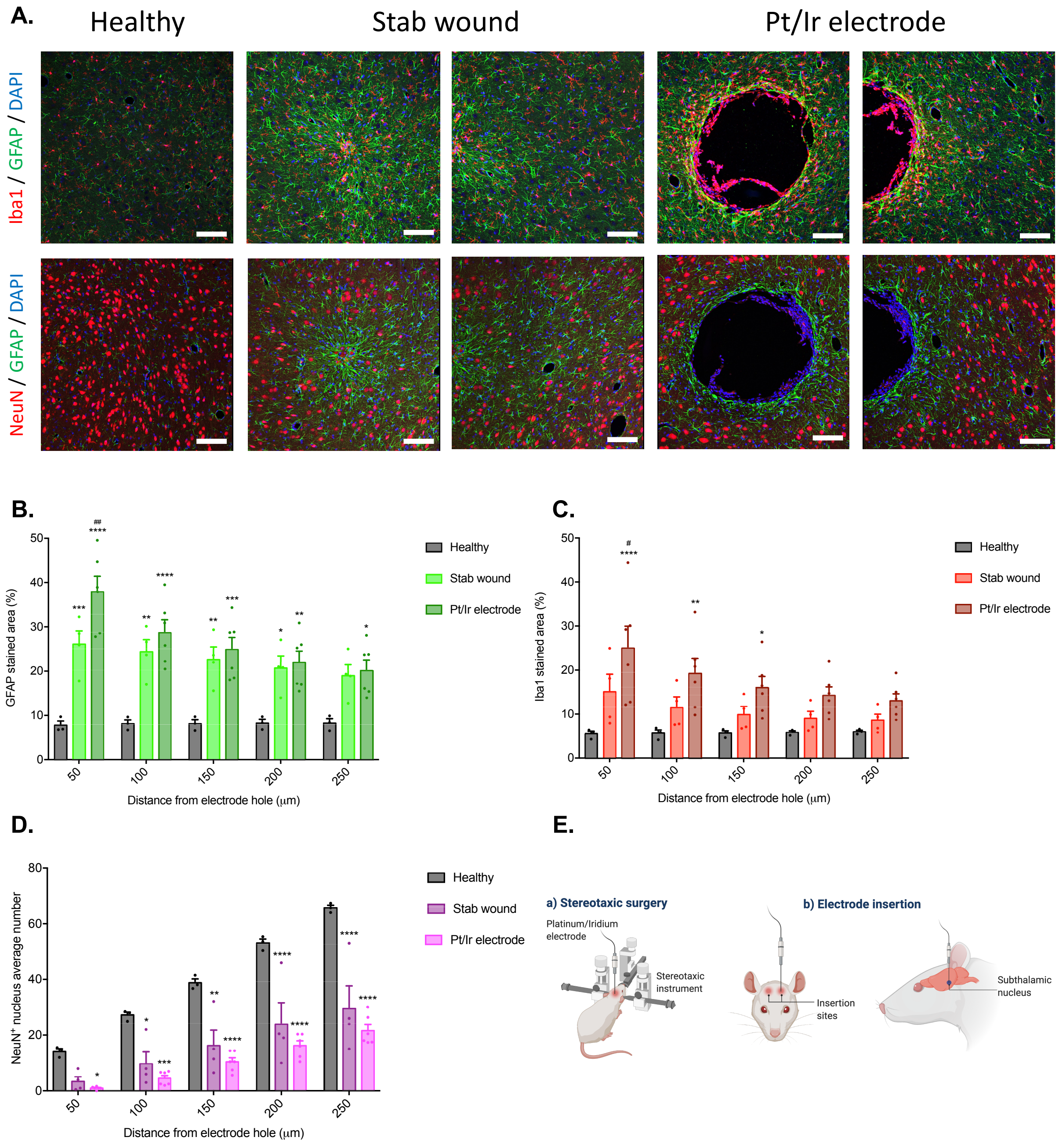


***Figure S7 Electrode implantation and stab wound injury lead to the glial scar development in vivo.*** *To confirm that the previously established in vivo model using stab injury or Pt/Ir electrode implantation into rat subthalamic nucleus for 8 weeks gave rise to the main hallmarks of the glial scarring process, immunofluorescent stainings of GFAP (green), Iba1 (red), NeuN (red) and DAPI (blue) were performed on the processed brain tissues (scale bar = 100µm)* ***(A)****. The GFAP stained area quantification of both the stab injury and implanted condition reveals a significant increase from ~58% to ~79% in comparison to the healthy control condition in function of the implantation site distance* ***(B)****. Similarly, the quantification of the Iba1 staining area percentage of the Pt/Ir electrode group in comparison to the healthy control condition manifested a significant increase from ~65% to ~78% as far as 150µm from the electrode hole* ***(C)****. Inversely, the number of mature neuron nucleus (NeuN$^{+}$) was significantly reduced from ~67% to ~83% in both the stabbed or implanted experimental groups when compared to the healthy control condition, as close as 100µm from the injury site* ***(D)****. Schematic detailing the insertion sites, created with BioRender.com* ***(E)****. Data are represented as mean ± SEM (n=3-6). Two-way ANOVA with Tukey post hoc test was performed. *, **, *** represents a statically significant difference versus the healthy control and #, ##, ### versus the stab wound condition, ($p<0.05$), ($p<0.01$) and ($p<0.001$), respectively.*

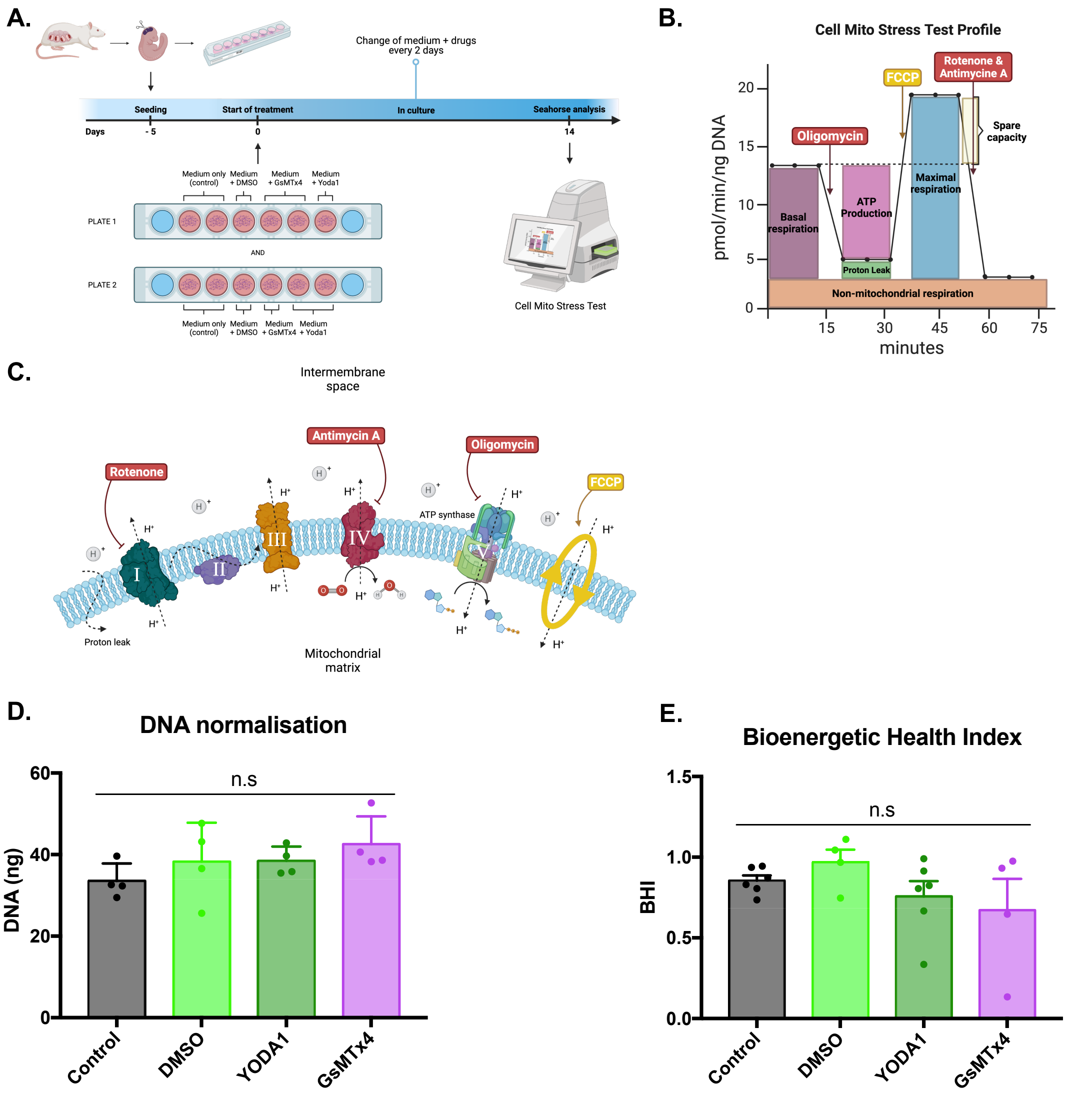


***Figure S8 Effect of the PIEZO1 receptor antagonist (GsMTx4) and agonist (Yoda1) on the mitochondria health state.*** *Schematic of the experiment design and timeline **(A)**. VM cells were extracted and seeded in Seahorse XF cell culture plates, after 5 days of culture growth and differentiation, cells were exposed to either medium only (control), medium and DMSO, medium and GsMTx4 or medium and Yoda1 and kept in culture for 14 days with a medium and drug refreshment every 2 days. Fourteen days after the start of the treatment, the Cell Mito Stress Test was performed using the Seahorse XF flux analyser. Graph representing the typical oxygen consumption rate (OCR) profile of healthy cells during the Mito Stress test **(B)**, the different mitochondrial functions (Basal respiration, ATP production, Maximal respiration, Spare capacity, Non-mitochondrial respiration and Proton leak) which can be directly calculated from these readings are displayed as well as the injection time of the different assay molecules. The different effects of the 4 molecules used in the Mito Stress Test **(C)** are pictured in this schematic **(C)**. The first injection, oligomycin is an inhibitor of the ATP synthase (complex V) which creates a drop in ATP production by reducing the electron flow through Electron Chain Transport (ETC), leading to a decreased mitochondrial respiration (OCR), this difference of respiration level correspond to the "ATP production" value. On the contrary, the second injection, Carbonyl cyanide-4 (trifluoromethoxy) phenylhydrazone (FCCP), an uncoupling agent, re-established the ETC by collapsing the proton gradient and disrupting the mitochondrial membrane potential which results to the highest mitochondrial OCR, value which correspond to the "Maximal respiration", moreover the difference between the basal respiration and maximal respiration allows to calculate the "Spare capacity" value which correspond to the mitochondria ability to respond to an increase metabolic demand. The last combined injection of Rotenone and Antimycine A, inhibitor of the complex I and III, respectively, stops entirely the mitochondrial respiration which allows to observe the "non-mitochondrial respiration" value, as well as the "proton leak", OCR difference between the full respiration inhibition and the sole complex V inhibition. All graph and schematics were created with BioRender.com. All flux analysis were normalized by the DNA amount (ng) of each well, DNA quantification which displayed similar values across all groups **(D)**. The bioenergetic health index calculation, indicator of the mitochondria health, showed no significant differences between both control and experimental groups **(E)**. Schematics created with BioRender.com.*
